# SNMF: Integrated Learning of Mutational Signatures and Prediction of DNA Repair Deficiencies

**DOI:** 10.1101/2024.11.27.624656

**Authors:** Sander Goossens, Yasin I. Tepeli, Colm Seale, Joana P. Gonçalves

## Abstract

**Motivation:** Many tumours show deficiencies in DNA damage response (DDR), which influence tumorigenesis and progression, but also expose vulnerabilities with therapeutic potential. Assessing which patients might benefit from DDR-targeting therapy requires knowledge of tumour DDR deficiency status, with mutational signatures reportedly better predictors than loss of function mutations in select genes. However, signatures are identified independently using unsupervised learning, and therefore not optimised to distinguish between different pathway or gene deficiencies.

**Results:** We propose SNMF, a supervised non-negative matrix factorisation that jointly optimises the learning of signatures: (1) shared across samples, and (2) predictive of DDR deficiency. We applied SNMF to mutation profiles of human induced pluripotent cell lines carrying gene knockouts linked to three DDR pathways. The SNMF model achieved high accuracy (0.971) and learned more complete signatures of the DDR status of a sample, further discerning distinct mechanisms within a pathway. Cell line SNMF signatures recapitulated tumour-derived COSMIC signatures and predicted DDR pathway deficiency of TCGA tumours with high recall, suggesting that SNMF-like models can leverage libraries of induced DDR deficiencies to decipher intricate DDR signatures underlying patient tumours.

**Availability:** https://github.com/joanagoncalveslab/SNMF.

## 1. Introduction

The accumulation of mutations caused by various DNA damage-inducing processes and associated DNA damage response (DDR) can lead to genome instability: an enabling hallmark of cancer (Hanahan and Weinberg, 2011) which can also be responsible for DDR pathway deficiencies frequently occurring in tumour cells.

Tumour DDR or repair deficiencies provide opportunities for targeted treatment, namely by leveraging known synthetic lethalities where joint disruption of multiple genes leads to cell death but individual inactivation does not affect cell viability (Kaelin, 2005). For example, patients with *BRCA1/2* deficient tumours can benefit from *PARP* inhibitor therapy targeting the synthetic lethal partner gene *PARP1*.

To apply treatments targeting DDR deficiency, it is necessary to identify which deficiencies might be present in the tumour. This could be done by detecting loss of function mutations in DDR related genes. However, this approach has limited sensitivity since not all genes involved in every repair pathway are known, and gene function can be modulated via alternative mechanisms such as epigenetic regulation (Yang *et al*., 2001). To capture downstream effects of alterations to DDR gene function, two models have been developed that predict deficiency in a specific repair pathway based on a number of features, including exposure to mutational signatures representing the patterns of mutations that such deficiency could leave on the genomes of tumour cells (Davies *et al*., 2017; Zou *et al*., 2021). Both models, HRDetect and MMRDetect, rely on a preselected subset of signatures from the ‘Catalogue of Somatic Mutations in Cancer’ (COSMIC) or similar, which are deemed relevant for the DDR deficiency of interest (Alexandrov *et al*., 2020). Signature identification for COSMIC is performed in an unsupervised manner, typically using non-negative matrix factorisation (NMF) (Alexandrov *et al*., 2013), and independently from the learning of a DDR deficiency prediction model. This means that the learned signatures are not optimised to discriminate between repair deficiency and proficiency, or deficiencies in different pathways. As a result, COSMIC contains multiple and partially redundant signatures associated with a given repair deficiency (e.g. 7 for mismatch repair deficiency) and many COSMIC signatures have unknown etiology. Since the mutational processes underlying each signature are not always evident, preselection of relevant DDR signatures becomes a challenge. Moreover, tumour samples do not have a ground truth repair deficiency status, and relying on proxy DDR deficiency labels based on loss of function mutations to DDR genes could lead to suboptimal prediction models.

We reason that supervised signature learning would be able to exploit knowledge of repair deficiency status to identify signatures that both (i) effectively distinguish DNA repair deficiencies, and (ii) capture more complete and representative signatures of each repair deficiency. Trained on mutational profiles with reliable DDR deficiency labels, such as experimental cell line data with induced DDR gene knockouts, supervised signature learning could not only lead to more robust models but also eliminate the need for manual annotation and preselection of signatures. On the other hand, cell lines are model systems that do not necessarily recapitulate what happens in tumours. Deeper investigation into the potential of supervised signature learning from cell line data is thus necessary. Supervised NMF methods have been proposed in other fields (Chao *et al*., 2019) that couple the NMF reconstruction to a classification objective, for instance by adding Fisher discriminant constraints (Wang *et al*., 2004), a Frobenius loss (Jing *et al*., 2012), or a cross-entropy loss (Serizel *et al*., 2017). The latter is relevant to our work, however we could not find the corresponding implementation or associated mathematical formulas. One other method, supervised negative binomial NMF (SNBNMF), was developed for mutational signature learning integrated with a support vector machine (SVM) to predict cancer subtype (Lyu *et al*., 2020). The SNBNMF models raw mutation counts using a negative binomial (NB) distribution. However, mutation counts can vary widely across samples, especially across domains such as between cell lines and tumours, genomes and exomes, or different cancer types. These differences could impair prediction generalisation across domains. To mitigate their impact, we prefer to use normalised counts, which do not follow the NB distribution. Here we propose Supervised NMF (SNMF), which integrates mutational signature learning and multiclass DDR deficiency prediction into a single model using a weighted loss with two terms: Frobenius reconstruction from NMF, and cross-entropy from multinomial logistic regression. We derive update formulas to optimise the SNMF model using gradient descentand evaluate SNMF against benchmark models. Finally, we apply SNMF to mutation profiles of human induced pluripotent stem (hiPS) cell lines harbouring knockouts of DDR-related genes, and investigate performance on patient tumour samples.

## 2. Methods

### 2.1. The SNMF Model

#### Problem formulation

##### Definitions

We define a matrix of mutation profiles ***X*** ∈ [0, 1]^*N×T*^, with *N* and *T* as numbers of samples and mutation types, respectively. An entry *x*_*n,t*_ represents the relative frequency of mutation type *t* in sample *n*, and each mutation profile (row in ***X***) is a probability distribution over all mutation types: 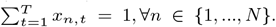 We also define the respective matrix of DNA repair deficiency status **Y** ∈ {0, 1}^N ×O^, over the *O* different repair deficiencies or classes the model can predict. The *n*^*th*^ row in ***Y*** represents the one-hot encoded repair deficiency status of sample *n*, where 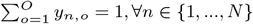.

##### Problem

Given a matrix of mutation profiles ***X***, the respective matrix of DNA repair deficiencies ***Y***, and a number of signatures *K* ≤ min(*T, N*), SNMF iteratively optimises: (i) mutation profiles in ***X*** as linear combinations of *K* underlying mutational signatures (matrix ***S***) weighed by exposures (matrix ***E***), and (ii) a linear mapping between sample exposures ***E*** and the output class or DNA repair deficiency ***Y*** with learned weights ***W***. Once built, the SNMF model can be used to estimate signature exposures and predict repair deficiency for mutation profiles of unseen samples.

#### Loss function

##### Reconstruction loss

To identify mutational signatures, matrix ***X*** is decomposed as a product of matrices ***S*** and ***E, X*** ≈ ***ES***. The signatures matrix ***S*** ∈ [0, 1]^*K×T*^ contains the relative frequencies of each of the *T* mutation types for each of the *K* signatures identified across samples, whereas the exposure matrix ***E*** ∈ [0, 1]^*N×K*^ contains the contribution of each signature to each of the *N* samples. We use NMF to find ***S*** and ***E*** (Lee and Seung, 2000), with multiplicative updates to optimise the Frobenius reconstruction error (*ℒ*_*r*_, Supplementary A.3).

##### Classification loss

For prediction of DNA repair deficiency, SNMF learns a linear mapping between signature exposures ***E*** and repair deficiency (class label) ***Y*** using multinomial logistic regression. This procedure estimates a matrix of weights ***W*** ∈ **R**^*K×O*^, with entry *w*_*k,o*_ denoting the weight of signature *k* with respect to the repair deficiency output class *o*. This is done by minimising the categorical cross-entropy loss (*ℒ*_*c*_) between observed (*y*) and predicted (*ŷ*) status for any given sample. To mitigate overfitting, we include L2 (ridge) regularisation, whose strength is controlled by hyperparameter *λ*_*L*2_ (Supplementary A.4).

##### Integrated (total) loss

To integrate both aims, SNMF optimises a combined loss (eq. 1), defined as the sum of the reconstruction *ℒ*_*r*_) and classification *ℒ*_*c*_) losses. The hyperparameter *λ*_*c*_ controls the integration between the two components: the larger the value, the larger the influence of the classification loss on exposures and signatures, while *λ*_*c*_ = 0 results in non-integrated signature decomposition (NMF) and repair deficiency prediction.

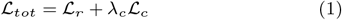

##### Multiplicative updates

The total loss *ℒ*_*tot*_ is minimised by iteratively applying gradient descent (GD) on *ℒ*_*tot*_ with respect to ***S, E***, and ***W***. The final updates are obtained by substituting the derivatives of the loss and the learning rates in the GD update formulas (⊙ and division are element-wise) (Supplementary A.1).

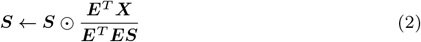

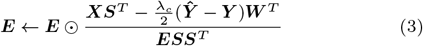

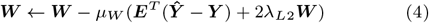

##### Non-negativity

The SNMF exposure update does not ensure non-negativity as a result of the subtraction between predicted and true labels (***Ŷ* − *Y***) (eq. 3). To ensure non-negativity, at each exposure update, negative values are set to a small positive value 10^−25^. We avoid using exactly 0 to prevent division by zero in the signature update (eq. 2). Each updated signature is normalised to keep it as a probability distribution over the *T* mutation types.

### 2.2. SNMF Model Learning and Prediction

We implement SNMF as an extension of SigProfiler (Alexandrov *et al*., 2013), offering regular NMF and a method to determine the optimal number of signatures. The application of SNMF comprises training (Fig. 1B-C) and testing (Fig. 1D), with input consisting of mutation profiles (and labels) of train and test samples (***X, Y, X***_***test***_), and hyperparameters (*K, λ*_*c*_, and *λ*_*L*2_).

**Fig 1:**
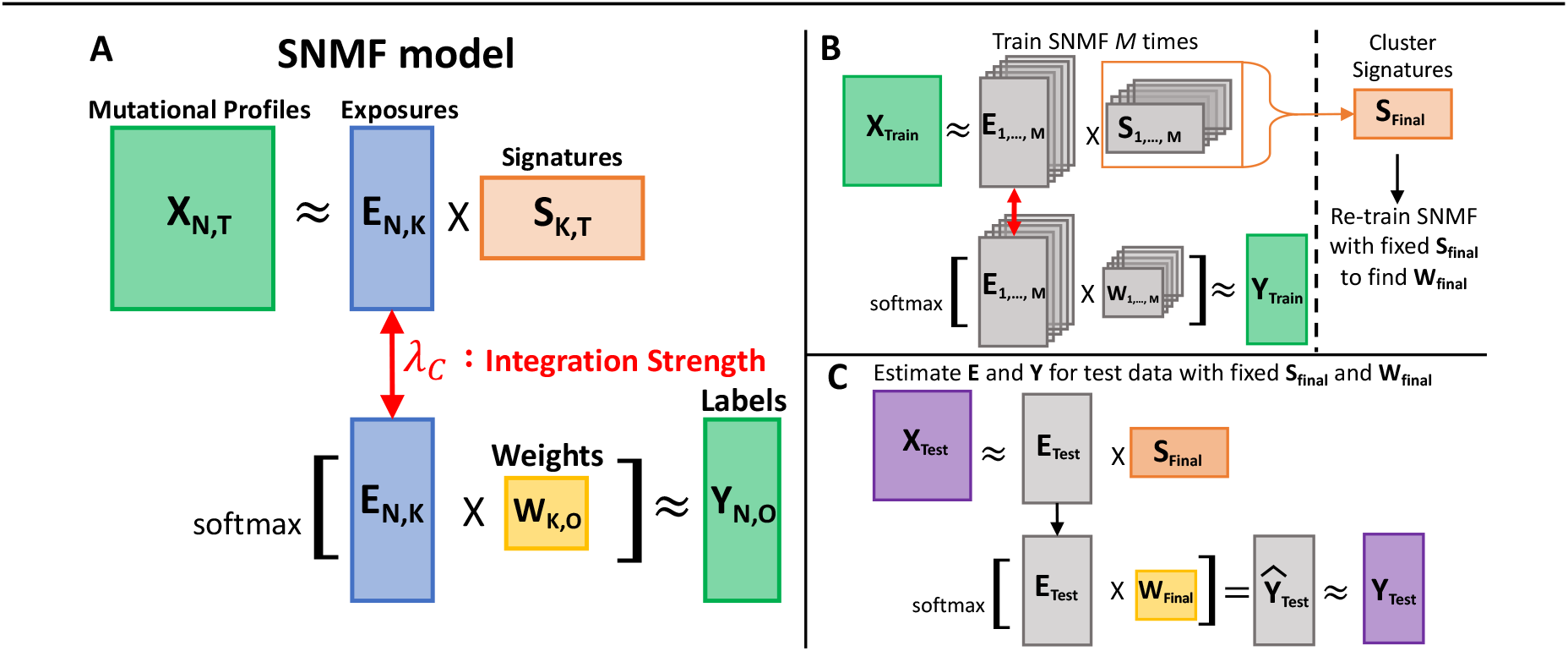
Overview of SNMF. **(A)** The SNMF model integrates (top) decomposition of mutation profiles into exposures and mutational signatures using NMF (***X*** ≈ ***ES***) and (bottom) exposure-based prediction of DNA repair deficiency using logistic regression (softmax(***EW***) ≈ ***Y***). **(B-D)** SNMF training and testing: (B) train *M* times to learn robust signatures ***S***_***final***_, (C) retrain with fixed ***S***_***final***_ to get regression weights ***W***_***final***_, and (D) obtain SNMF exposures and predictions for test data with fixed ***S***_***final***_ and ***W***_***final***_.

The SNMF model is trained by randomly initialising ***S*** and ***E***, and then iteratively updating ***S, E*** and ***W*** (eq. 2-4), resulting in a set of *K* signatures with respective exposures and regression weights. To account for the variation caused by random initialisation, training is repeated *M* times, after which the *M × K* signatures found across runs are clustered using a variation of K-means (Fig. 1B), where each cluster is assigned one signature from each run. The final set of *K* signatures ***S***_***final***_ is obtained by averaging the signatures within each cluster. The signatures ***S***_***final***_ are then fixed and fitted to obtain the exposures ***E*** and weights ***W***_***final***_ (Fig. 1B) using non-integrated SNMF (*λ*_*c*_ = 0).

The signatures ***S***_***final***_ are fixed and fitted to the unseen test samples ***X***_***test***_ using non-negative least squares (NNLS) to obtain the respective exposures ***E***_***test***_ (Fig. 1C). The test exposures ***E***_***test***_, together with the classification weights ***W***_***final***_ of the trained SNMF model, are used to calculate the prediction probability of each deficiency class for every test sample (***Ŷ***). The final predicted label is the class with the highest probability.

### 2.3. Data and Preprocessing

#### Cell line mutation profiles

We trained SNMF using data from Zou *et al*. (2021), profiling mutations for hiPS cell lines with biallelic knockouts (KOs) of 42 different genes involved in 12 DNA repair pathways. Knockout of gene *ATP2B4* was also included as a control. There were 2 to 8 replicates per gene KO, resulting in 173 samples. Mutations were obtained after culturing for 15 days, whole-genome sequencing, alignment to the human reference GRCh37/hg19, and somatic mutation calling. We focused on 96 types of single base substitutions (SBS) comprising: the substitution itself, with 6 different types considering pyrimidines C and T as reference (G and A are complementary); and the two neighbouring 5’ and 3’ bases, resulting in 16 possible for each SBS. The original count-based mutation profiles were normalised to represent a probability distribution over the 96 SBS types. In line with Zou *et al*. (2021), we selected gene KOs for which all replicates had either high mutation count and/or low cosine similarity with the average profile of control samples (Supplementary Fig. S2).

#### Cancer patient mutation profiles and signatures

Cell line signatures found by SNMF were compared to 78 cancer-related COSMIC signatures (v3.2). To assess SNMF prediction for patient tumours, we used whole-exome MC3 somatic mutations from The Cancer Genome Atlas (TCGA, Ellrott *et al*. (2018)). For HR deficiency (HRd) prediction, we focused on breast cancer (BRCA), since HRd is most prevalent in BRCA and ovarian (OV) cancers (Venkitaraman, 2014), with a much larger number of BRCA tumours (791 vs. 65 OV). For MMR deficiency (MMRd), we evaluated SNMF prediction on the three tumour types with highest prevalence of microsatellite instability (MSI): uterine (UCEC), colon (COAD), and stomach (STAD) (Bonneville *et al*., 2017). We used repair status of patient tumours based on mutation status of 81 repair genes assigned to 9 pathways (Knijnenburg *et al*., 2018). Each was annotated as wild-type, or heterozygous or homozygous mutated based on variant allele frequency of either loss of function mutations or epigenetic silencing.

### 2.4. Data Augmentation and Hyperparameter Optimisation

To build the SNMF model, we split the cell line data into disjoint train and test sets. The test set contained exactly one replicate of each gene KO and 2 replicates of the control samples. The remaining train samples were split into three folds for cross-validated hyperparameter optimisation, with replicates per gene KO evenly spread across folds (Supplementary Fig. S3).

#### Data augmentation

To improve SNMF model robustness, given the limited number of cell line samples, we augmented the data using bootstrapped oversampling. Per fold, we oversampled the number of train samples to 100 per class using the real gene KOs. Each new bootstrapped sample was created by considering the mutation profile of a real sample as a probability distribution, based on which we drew the same number of mutations as in the real sample with replacement. The counts of the new bootstrapped samples were divided by the sum so each mutation profile summed to 1. For testing, we bootstrapped up to 1000 samples per class, since prediction is more efficient than model learning.

#### Hyperparameter optimisation

The SNMF model has as parameters: number of signatures *K*, integration strength *λ*_*c*_, and regularisation strength *λ*_*L*2_. Since the number of signatures had the most impact on performance, we first determined *K* as the value that resulted in the highest accuracy without a decrease in stability across a range of *λ*_*c*_ and *λ*_*L*2_. Then we selected *λ*_*c*_ and *λ*_*L*2_ together based on the optimal Pareto settings, containing all parameter settings for which no other yielded higher accuracy and stability. Using the optimal setting, we trained the final integrated SNMF model on the entire cell line train set.

### 2.5. Evaluation of the SNMF Model

The cell line-trained SNMF model was first evaluated and compared to benchmark models on the cell line test samples.

#### Evaluation metrics

We used three metrics: stability of identified signatures, and both prediction accuracy and reconstruction error on test samples. Signature stability or reproducibility was measured using the average of silhouette widths (ASW) of the *K* signature clusters (*Cluster*_1_, …, *Cluster*_*K*_) obtained with *M* training runs, based on cosine distance between signatures. The (average) SW ranges between -1 and 1, with 1 indicating consistent signature(s) across runs and lower SW values denoting weaker reproducibility. We used accuracy to assess classification performance, corresponding to the fraction of correctly predicted test samples (true positives and true negatives), out of all test samples. The reconstruction error evaluated how well the mutation profiles of the test samples could be recovered from the exposures and (fixed) signatures, defined based on the Frobenius norm as follows: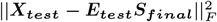. The lower the error, the more accurate the approximation to the original test data ***X***_***test***_. Since choosing a larger number of signatures *K* typically results in lower reconstruction error, owing to the higher model complexity, we considered the reconstruction subordinate to stability for measuring the quality of signature decomposition.

#### Benchmark models

We assessed the integrated SNMF model with joint signature learning and repair deficiency prediction (*λ*_*c*_ *>* 0) against two benchmarks: 1) an L2-regularised logistic regression model learned directly from the mutation profiles ***X***, thus using mutation profiles rather than signature exposures as input; and 2) non-integrated signature learning and exposure-based prediction using SNMF with *λ*_*c*_ = 0, equivalent to existing approaches that first apply unsupervised NMF signature learning and then build an exposure-based classification model.

### 2.6. Exposures and Predictions for TCGA Patient Tumours

The final SNMF model was used to obtain exposures and make predictions for TCGA patient tumour samples. We used recall to assess the ability to identify repair deficiencies in tumours with a homozygous mutated core gene for the repair pathway of interest. We preferred recall to accuracy, since the absence of loss of function mutations in a repair gene does not imply repair proficiency.

Given that genes involved in a repair pathway have different functions and could lead to distinct repair deficiency signatures, we also analysed the SNMF results for patient tumours on a gene rather than pathway level. To quantify the association between the mutation status of a repair gene and exposure to the corresponding SNMF deficiency signature, we performed a one sided Mann-Whitney U (MWU) test between the exposures (*E*_*MMRd*_ or *E*_*HRd*_) of mutated and non-mutated tumours, resulting respectively in statistics U1 and U2 together with the MW p-value. We did not distinguish heterozygous and homozygous mutations, as very few tumours carried homozygous mutations for most genes. We also quantified the co-occurrence of mutations for each pair of genes using Jaccard similarity.

## 3. Results and Discussion

### 3.1. Mutation Profiles and Repair Deficiencies

Nine gene KOs showed mutation profiles distinguishable from the control samples (Fig. 2A), and were annotated with a total of four repair pathways: mismatch repair (MMR: *MSH6, MSH2, MLH1, PMS2* and *PMS1*), base excision repair (BER: *UNG* and *OGG1*), homologous recombination (HR: *EXO1*), and double strand break repair (DSB: *RNF168*). The high similarity of *RNF168* and *EXO1* (cossim: 0.96) and literature supporting the role of *RNF168* in recruiting the *BRCA1-PALB2* complex to DNA damage for HR-directed repair (Krais *et al*., 2021), led us to annotate *RNF168* with HR as well. Resulting in four classes: MMRd, BERd, HRd, and control (*ATP2B4*). As expected, bootstrapped samples clustered around the gene KO they were sampled from (Fig. 2B). We also confirmed that *RNF168* and *EXO1*, indeed clustered together. On the other hand, *OGG1* and *UNG* formed their own clusters, indicative of distinct behaviour within BER. In addition, we observed that *PMS1* was not as discernable from the control samples as the other MMR gene KOs. Additionally, *PMS2* did not cluster together with the other MMR gene KOs, yet it was distinctive from the control samples.

**Fig 2:**
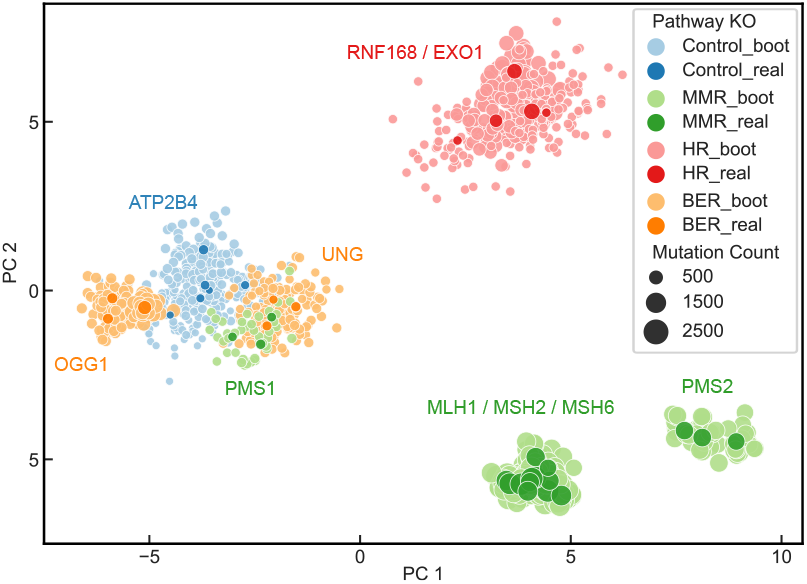
Gene knockout and bootstrapped training samples. Latent PCA plot of bootstrapped (light) and real (dark) tumour samples used for training.

### 3.2. Influence of SNMF Hyperparameters

The number of signatures *K* had the largest effect on accuracy, stability, and reconstruction (Fig. 3A). Increasing *K* from 3 to 7 had a positive effect on median accuracy (0.78 vs 0.93) and reconstruction error (5.74 vs 4.57). This was expected, since using more signatures increases model complexity and could result in a better fit to the data. This effect was less pronounced above 5 signatures. The median stability improved when increasing *K* from 3 to 5 signatures (0.67 vs 0.99), but decreased for *K >* 5 (0.72 for *K* = 7). This was also apparent in the Pareto plot (Fig. 3B, green vs red), where extracting 7 signatures with optimal integration and regularisation strengths resulted in higher accuracy compared to 5 signatures (0.964 vs 0.942), at the cost of stability (0.46 vs 1.0). The likely cause is that, for 7 signatures, SNMF occasionally found a signature specific to *PMS1* gene KOs (Fig. 3C, red), but not consistently across runs. We settled on 5 signatures (*K* = 5), since it resulted in the most reproducible signatures.

**Fig 3:**
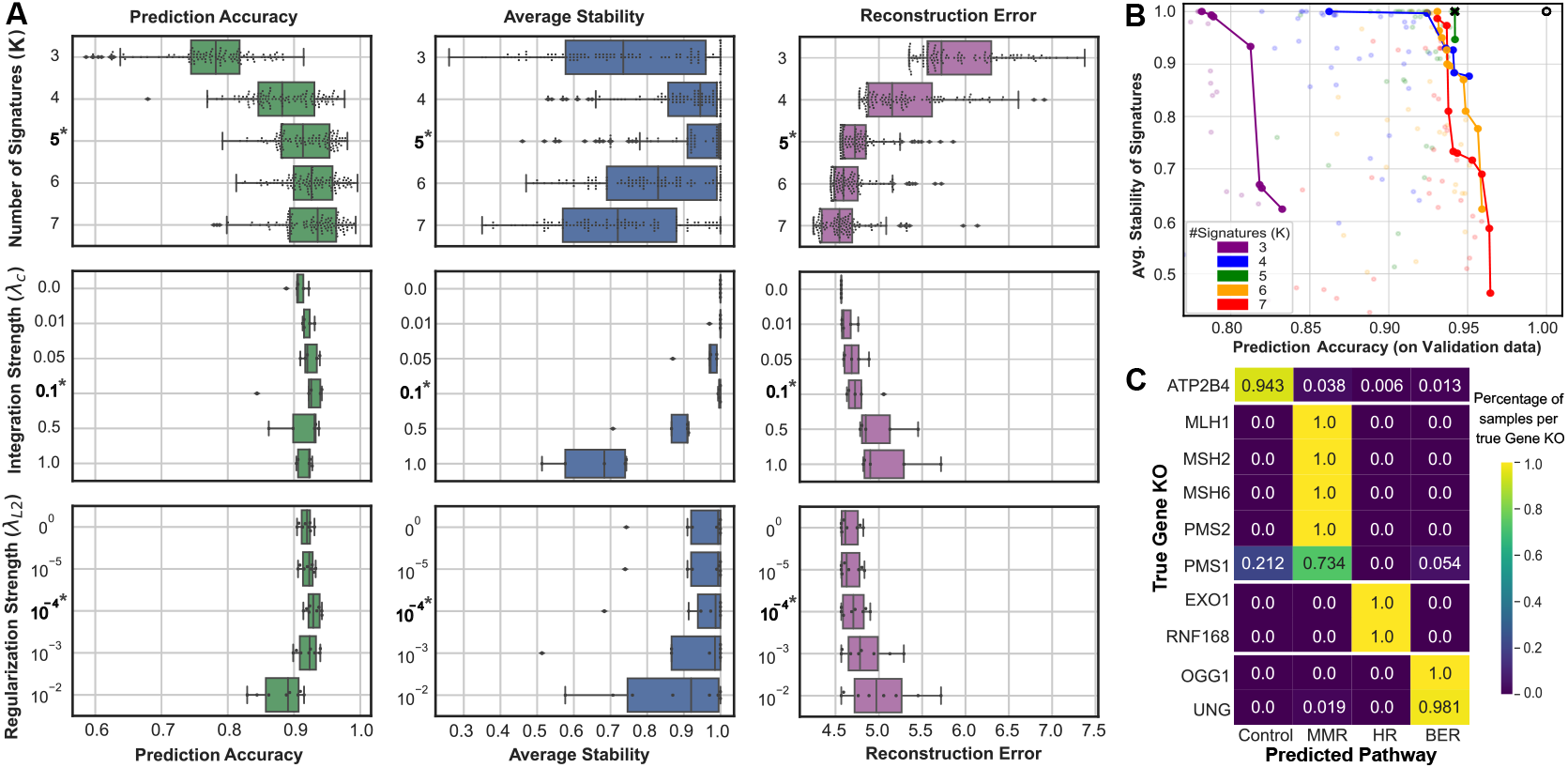
Hyperparameter optimisation and prediction performance of SNMF. **(A)** Boxplots of hyperparameter optimisation results using cross-validation. Each dot denotes the average evaluation metric (Left: accuracy; Middle: stability; Bottom: reconstruction) of three validation folds per hyperparameter setting: (Top) number of signatures *K*; (Middle row) integration *λ*_*C*_ (with *K* = 5); and (Bottom) regularisation *λ*_*L*2_ (with *K* = 5). Final settings indicated in bold (*\**) **(B)** Optimal *λ*_*C*_ and *λ*_*L*2_ based on accuracy and stability per number of signatures *K*. Each dot denotes an average over three validation folds for one hyperparameter setting. Lines indicate pareto optimal *λ*_*C*_ and *λ*_*L*2_ per *K*. (***×***, final optimal setting) (○, theoretical perfect setting). **(C)** Confusion matrix of SNMF predictions per gene KO. Coloured and annotated by percentage of samples per gene KO (real and bootstrapped) predicted to each pathway.

We looked into the impact of integration and regularisation strengths using *K* = 5 signatures. Higher integration strength improved prediction accuracy, from median 0.906 with no integration (*λ*_*c*_ = 0.0) to median 0.931 with integration (*λ*_*c*_ = 0.5), with high stability. However, further increasing the integration strength deteriorated both stability and reconstruction error, owing to a dominance of prediction over signature identification.

Using a small regularisation strength slightly increased the median accuracy from 0.920 without regularisation to 0.938 with *λ*_*L*2_ = 0.0001. Too strong regularisation not only led to a decrease in accuracy (median 0.899 for *λ*_*L*2_ = 0.01), it also worsened stability and reconstruction error. This highlights the integrated nature of the SNMF model, where the direct effect of regularisation on classifier weights also indirectly affects signature learning.

We analysed the tradeoff between accuracy and stability for various hyperparameter settings using Pareto plots (Fig. 3B). The optimal settings form a front per *K*, where the best performing is closest to the theoretically perfect model with accuracy and stability equal to 1.0 (Fig. 3B, black *○*). For *K* = 5, the optimal setting yielded integration *λ*_*c*_ = 0.1 and regularisation *λ*_*L*2_ = 0.0001 (Fig. 3A-3B black *×*), with accuracy 0.942 and stability 1.0. These settings were used to train the final SNMF model.

### 3.3. Integrated SNMF Has High Prediction Accuracy

Integrated SNMF achieved high prediction accuracy (0.971), comparable to non-integrated SNMF (0.966) and logistic regression (0.968). This was accompanied by good mutational signature decomposition performance, similar to unsupervised NMF (stability: 0.99 vs 1.0 and reconstruction error 4.57 vs 4.38, Table 1). The logistic regression model, being prediction-centric and without signature discovery, was thus not as easily relatable to the underlying biological processes as the SNMF model.

**Table 1.** Performance comparison of final models.

| Model | Accuracy | Stability | Reconstruction |
| --- | --- | --- | --- |
| Logistic regression | 0.968 | - | - |
| Non-integrated SNMF | 0.966 | **1.00** | **4.38** |
| **Integrated SNMF** | **0.971** | 0.99 | 4.57 |

For integrated SNMF, most incorrect predictions were made between control and MMRd samples (Fig. 3D), where 21.2% of the *PMS1* gene KO samples were misclassified as control (accuracy 0.739). This was in line with the clustering of *PMS1* KO samples closer to control samples than to other MMRd gene KOs (fig. 2). The remaining MMRd and HRd gene KO samples were predicted perfectly (1.0), and accuracy was also high for BERd (*>*0.98).

For both integrated and non-integrated SNMF, all real samples were predicted correctly, with misclassifications observed only for bootstrapped samples. The non-integrated SNMF showed a higher proportion of misclassifications between *PMS1* and control samples, but all remaining gene KOs were predicted with high accuracy (Supplementary Fig. S6). Finally, the logistic regression model mainly misclassified control samples as BERd, while most PMS1 samples were predicted correctly (0.94).

### 3.4. SNMF Identifies BER Subpathway Deficiencies

The SNMF model predicted all BERd samples with high accuracy, but also identified two signatures clearly distinguishing the *OGG1* and *UNG* gene knockouts (Fig. 4A, brown and green respectively). *OGG1* and *UNG* both encode for DNA glycosylase enzymes, but repair different types of DNA damage. UNG recognises and excises uracil (deaminated bases), and its deficiency leads to C*>*T mutations when the DNA is replicated; whereas OGG1 recognises and removes 8-oxoG (oxidised bases) and its deficiency results in G*>*T (or C*>*A) mutations (Supplementary Fig. S9) (Zou *et al*., 2021). The ability to recognise subpathways within a given deficiency class is a key advantage of an NMF-based model over the direct logistic regression, or any other prediction-focused model that does not aim to capture underlying mutational patterns.

**Fig 4:**
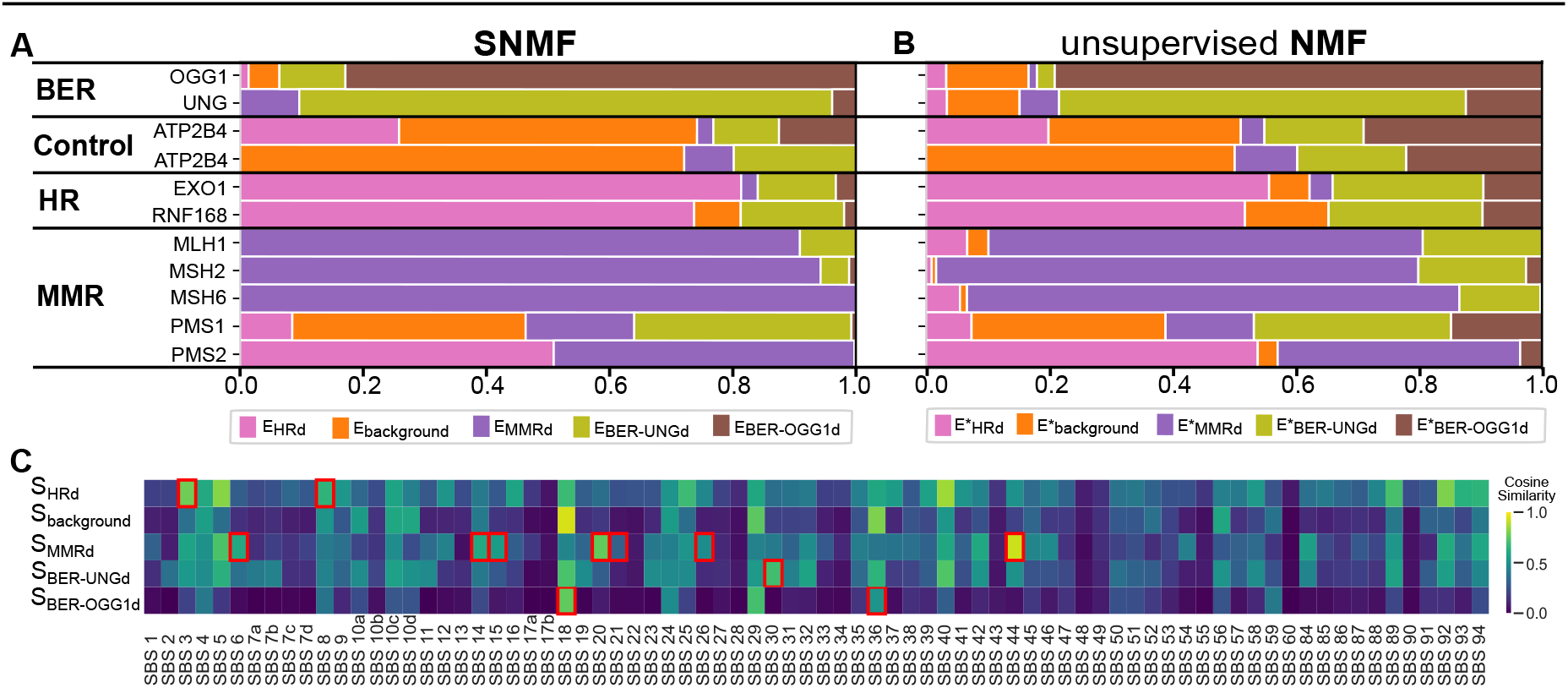
Cell line signatures identified by SNMF and NMF. **(A-B)** Signature exposures found by (A) SNMF and (B) NMF for the real test cell line samples (no bootstrapped samples). **(C)** Heatmap showing similarity between cell line signatures found by SNMF and COSMIC signatures. Red squares highlight COSMIC signatures with aetiology related to each DNA repair pathway deficiency.

### 3.5. SNMF Signatures Distinguish Repair Deficiencies

We compared unsupervised NMF and integrated SNMF by pairing signatures based on cosine similarity (Supplementary Fig. S9).

The SNMF model mainly used one signature to decompose each sample, with an average of the largest exposure per sample of 0.80 (Fig. 4A). In contrast, unsupervised NMF led to more mixed exposures to additional signatures, with a largest exposure of only 0.61 on average. In this context, one could be led to interpret the contribution of multiple signatures as an indication that cells have multiple repair deficiencies instead of one, such as suggested by the exposure of MMRd samples to the BER-UNG signature (Fig. 4B, green). The exposures of unrelated signatures were reduced using integrated SNMF compared to the NMF model, suggesting that each repair deficiency was better captured into one signature by exploiting the information contained in the class labels.

The unsupervised NMF tended to isolate a mutation type into one signature and to exclude it from other signatures. This effect was reduced using integrated SNMF, where multiple signatures included similar characteristic mutations (Supplementary Fig. S9).

### 3.6. Cell line Signatures Recapitulate Cancer Signatures

We analysed the cell line signatures found by integrated SNMF against cancer COSMIC signatures linked to the same repair deficiency (Fig. 4C). Of all COSMIC signatures, only SBS3 has been annotated with HRd. Davies *et al*. (2017) have also suggested that SBS8 could be linked to HRd, and use exposures of SBS8 and SBS3 as predictors in the classifier HRDetect. The HRd signature showed high similarity to SBS3 (cossim: 0.78) and SBS8 (cossim: 0.65). Seven partially overlapping COSMIC signatures are linked to MMRd (Alexandrov *et al*., 2020), which studies have suggested can be reduced to two main processes (Németh *et al*., 2020). One MMRd signature is characterised by T*>*C mutations, has high similarity to SBS21 and SBS26, and relates to *PMS2* -based MMRd; the other contains C*>*T mutations and has high similarity to SBS6, SBS15, and SBS44 and relates to *MLH1/MSH2/MSH6* -based MMRd (Supplementary Fig. S9). (Degasperi *et al*., 2022). The MMRd signature found by SNMF was most similar to the latter (SBS44), explaining why *PMS1/2* are not as well capture by the signature. For BERd, two COSMIC signatures relate to *OGG1* deficiency: SBS18 annotated with DNA damage by reactive oxygen, and SBS36 linked to *MUTYH* deficiency. The BER-OGG1d signatures found by SNMF showed high similarity to SBS18 (cossim: 0.76), and to a lesser extent SBS36 (cossim: 0.52). The BER-UNGd was most similar to SBS30 (cossim: 0.89), related to *NTHL1* deficiency, which, like *UNG*, is a BER glycosylases that recognize abnormal pyrimidines (Zou *et al*., 2021).

### 3.7. Cell line SNMF Predicts Tumour Repair Deficiencies

We evaluated the ability of SNMF trained on cell line data to generalise to tumours. All tumours with a homozygous mutated MMR gene were predicted as MMRd (recall 1.0), most carrying *MLH1* mutations (30*/*37). Additionally, eleven of the twelve tumours yielding a homozygous mutated HR gene were predicted as HRd (recall 0.92), where the misclassified tumour was BRCA1 deficient. The SNMF model was more sensitive to the exposure of the MMRd signature than to the exposure of the HRd signature for prediction of MMRd and HRd (Supplementary Fig. S14).

For the MMRd related cancer types, mutations in five genes (*MLH1, APEX2, FANCB, MSH2, MSH6*) significantly associated with *E*_*MMRd*_ (Fig. 5A). Mutations in the *MLH1* gene had a strong effect on *E*_*MMRd*_, confirming the known role of MLH1 in MMRd (Mensenkamp *et al*., 2014). The effect for APEX2 and FANCB was no longer significant after excluding tumours with co-occuring mutations in other MMR genes, and we did not find literature linking APEX2 or FANCB to MMR. The MMR genes *MSH2* and *MSH6*, mutated in a small number of tumours (28 and 32), initially associated with *E*_*MMRd*_ (*p* ≤ 0.05). But upon exclusion of tumours with mutations in other MMR genes, too few samples remained (9 and 6) to test the effect. Interestingly, tumours with high *E*_*MMRd*_ were depleted of *TP53* mutations.

**Fig 5:**
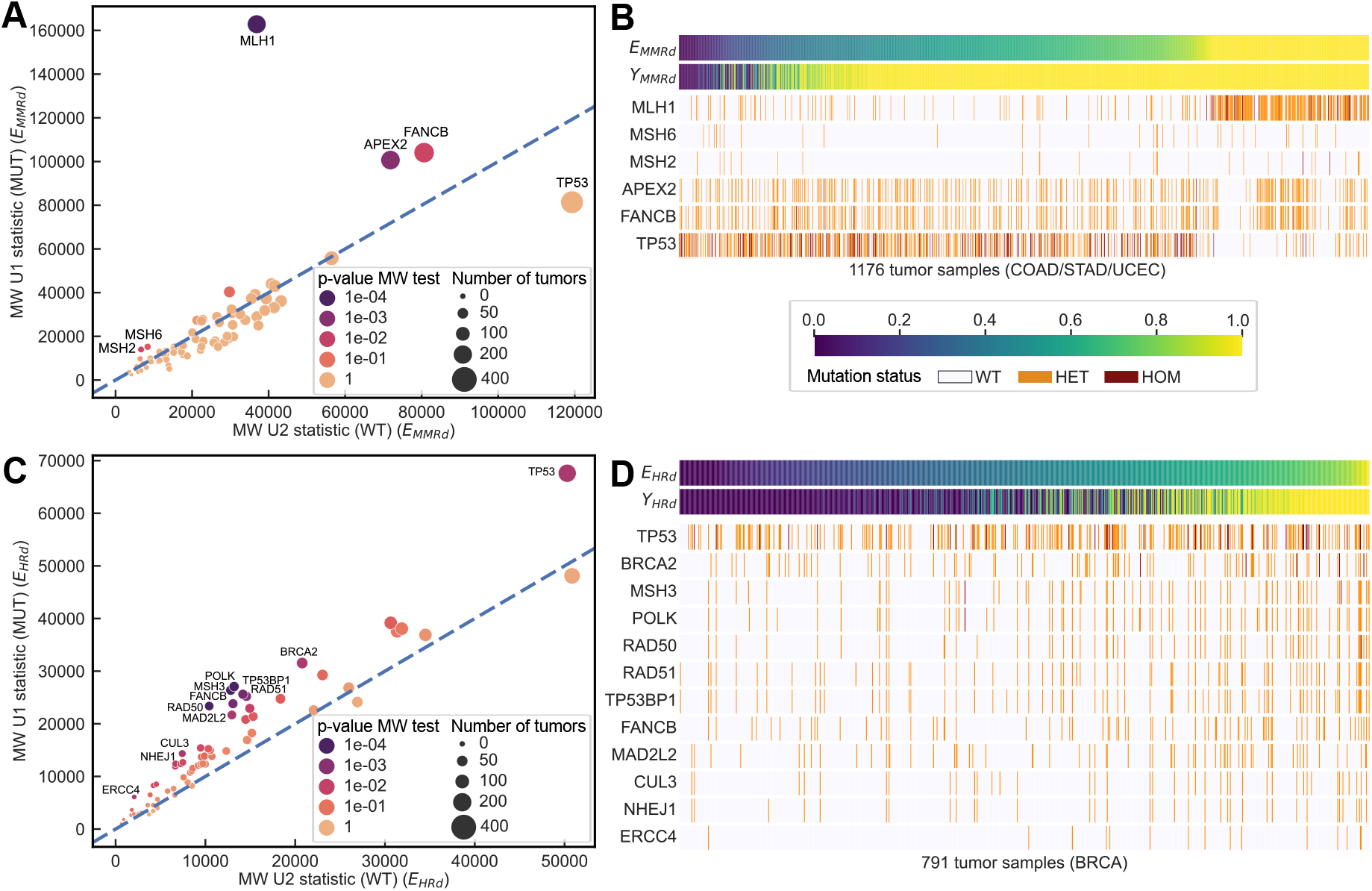
Integrated SNMF predictions on TCGA tumour data, and genes related to the MMR and HR repair deficiencies. **(Top, A-B)**: MMRd in COAD, STAD and UCEC tumours. **(Bottom, C-D)**: HRd in BRCA tumours. **Left:** Mann-Whitney U test statistics per repair gene, comparing the signature exposure for wild-type tumours (U2) against tumours with the mutated gene (U1)**Right:** Top two rows represent the exposure and prediction probability found by SNMF (**B**: *E*_*MMRd*_ and *Y*_*MMRd*_, **D**: *E*_*HRd*_ and *Y*_*HRd*_). The remaining rows represent mutation status of tumour samples for the genes with the lowest MWU p-value (**(B)** MMRd *p* ≤ 0.05, **(D)** HRd *p* ≤ 0.01). For MMRd, *TP53* is included to show negative correlation with *E*_*MMRd*_.

For HRd, 24 genes yielded an association with *E*_*HRd*_ (*p* ≤ 0.05), of which 12 were more strongly significant (*p* ≤ 0.01) and 8 still showed an effect after excluding tumours with mutations in other HR genes (Supplementary Fig. S16). Among this latter group were four core HR genes (*TP53BP1, BRCA2, RAD51*), two genes related to HR (*POLK, TP53*), and 4 genes linked to other processes (*MSH3, FANCB, MAD2L2*). The effect of FANCB on HR supports the known interplay between fanconi anemia (FA) and HR pathways, as well as the role of FANCB in the FA-BRAC pathway (Meetei *et al*., 2004). Other HR genes linked to increased risk of cancer, such as *BRCA1, PALB2, BARD1, BRIP1*, or *ATM* (Tsang *et al*., 2023), were not found more frequently mutated in tumours with high *E*_*HRd*_. It has been suggested that there could be two types of HRd, *BRCA1* -HRd and *BRCA2* -HRd, resulting in distinct mutational patterns (Nguyen *et al*., 2020). A potential explanation for the absence of a significant effect of *BRCA1* and *BRCA1* -related genes could be that the HRd signature found by SNMF (*S*_*HRd*_) does not capture *BRCA1* -HRd.

## 4. Conclusion

We introduced SNMF, a supervised NMF approach that integrates mutational signature learning with multinomial logistic regression for prediction of repair pathway deficiencies. The SNMF model achieved high accuracy for predicting repair deficiency of hiPS cell lines, with high signature stability and low reconstruction error. The number of signatures had the most impact on SNMF performance, and larger integration strength reportedly improved accuracy at the cost of signature stability. Attesting to the value of the supervised component, signatures extracted by SNMF provided a more complete representation of each repair deficiency class, whereas unsupervised NMF tended to split patterns of a deficiency into multiple signatures. Additionally, both (S)NMF captured signatures linked to distinct subpathways, such as OGG1 and UNG in BER. The cell line SNMF model further recapitulated tumour-related COSMIC signatures and identified repair deficiencies in TCGA tumours with high recall. However, the SNMF classification component was too sensitive to the MMR signature exposure. This could be improved in the future by allowing for a rejection option and adapting the model for prediction in the tumour domain, for instance using semi-supervised learning. Gene-level analysis suggested that deficiency in genes linked to a given pathway can lead to distinct mutation patterns, such as BRCA1-vs BRCA2-HRd, or MLH1-vs PMS2-MMRd, underlining the importance of discerning subtype repair deficiencies. Beyond the benefits of supervised signature learning for DDR prediction using SNMF, we show the potential of cell line SNMF signatures to unravel repair deficiencies in patient tumours. This creates opportunities for interpretable and clinically relevant prediction models aimed at improving precision cancer therapy.

## Acknowledgments

Authors received funding from: Convergence of Health and Technology [2022035 to J.P.G.]; and the US National Institutes of Health [U54DK134302 to S.G. and J.P.G.] and [U54EY032442, U01DK133766, R01AG078803 to J.P.G.]. Authors are solely responsible, funders were not involved in the work.

## References

Alexandrov, L. B., et al. (2013). Deciphering Signatures of Mutational Processes Operative in Human Cancer. Cell Rep., 3, 246–259.

Alexandrov, L. B., et al. (2020). The repertoire of mutational signatures in human cancer. Nature, 578(May 2018).

Bonneville, R., et al. (2017). Landscape of Microsatellite Instability Across 39 Cancer Types. Technical report.

Chao, G., et al. (2019). Recent Advances in Supervised Dimension Reduction: A Survey. Machine Learning and Knowledge Extraction, 1(1), 341–358.

Davies, H., et al. (2017). HRDetect is a predictor of BRCA1 and BRCA2 deficiency based on mutational signatures. Nature Medicine, 23(4), 517–525.

Degasperi, A., et al. (2022). Substitution mutational signatures in whole-genome–sequenced cancers in the UK population. Science, 376(6591).

Ellrott, K., et al. (2018). Scalable Open Science Approach for Mutation Calling of Tumor Exomes Using Multiple Genomic Pipelines. Cell Systems, 6(3), 271–281.

Hanahan, D. and Weinberg, R. A. (2011). Hallmarks of cancer: The next generation. Cell, 144(5), 646–674.

Jing, L., et al. (2012). SNMFCA: Supervised NMF-based image classification and annotation. IEEE Transactions on Image Processing, 21(11), 4508–4521.

Kaelin, W. G. (2005). The concept of synthetic lethality in the context of anticancer therapy. Nat. Rev. Cancer, 5, 689–698.

Knijnenburg, T. A., et al. (2018). Genomic and Molecular Landscape of DNA Damage Repair Deficiency across The Cancer Genome Atlas. Cell Reports, 23(1), 239–254.

Krais, J. J., et al. (2021). RNF168-mediated localization of BARD1 recruits the BRCA1-PALB2 complex to DNA damage. Nature Communications, 12(1).

Lee, D. D. and Seung, H. S. (2000). Learning the parts of objects by non-negative matrix factorization. Nature. Nature, 401(October 1999), 788–791.

Lyu, X., et al. (2020). Mutational signature learning with supervised negative binomial non-negative matrix factorization. Bioinformatics, 36, 154–160.

Meetei, A. R., et al. (2004). X-linked inheritance of Fanconi anemia complementation group B. Nat. Gen., 36, 1219–1224.

Mensenkamp, A. R., et al. (2014). Somatic mutations in MLH1 and MSH2 are a frequent cause of mismatch-repair deficiency in lynch syndrome-like tumors. Gastroenterology, 146, 643–646.

Németh, E., et al. (2020). Two main mutational processes operate in the absence of DNA mismatch repair. DNA Repair, 89(February), 102827.

Nguyen, L., et al. (2020). Pan-cancer landscape of homologous recombination deficiency. Nature Comm., 11(1), 1–12.

Serizel, R., et al. (2017). Supervised group nonnegative matrix factorisation with similarity constraints and applications to speaker identification. IEEE ICASSP, pages 36–40.

Tsang, E. S., et al. (2023). Homologous recombination deficiency signatures in gastrointestinal and thoracic cancers correlate with platinum therapy duration. npj Precision Oncology, 7(1).

Venkitaraman, A. R. (2014). Cancer suppression by the chromosome custodians, BRCA1 and BRCA2. Science, 343(6178), 1470–1475.

Wang, Y., et al. (2004). Fisher non-negative matrix factorization for learning local features. Proceedings of the Asian Conference on Computer Vision, pages 27–30.

Yang, X., et al. (2001). DNA Methylation in Breast Cancer. Endocrine-related Cancer, 8, 115–127.

Zou, X., et al. (2021). A systematic CRISPR screen defines mutational mechanisms underpinning signatures caused by replication errors and endogenous DNA damage. Nature Cancer, 2(6), 643–657.

