## Supplementary Material for "SNMF: Integrated Learning of Mutational Signatures and Prediction of DNA Repair Deficiencies"

A. Extended Methods: derivation of the update rules and derivatives for the S-NMF optimization algorithm

#### A.1. Matrix definitions

$\boldsymbol{X}_{N,T}$ : Input matrix of (normalized) frequencies of $T$ mutation types for each of the $N$ samples (mutational profiles)

$\boldsymbol{Y}_{N,O}$ : Output class label matrix, with $O$ class labels for each of the $N$ samples (one-hot encoded DNA repair deficiencies)

$\boldsymbol{S}_{K,T}$ : Mutational signature matrix, denoting the frequencies of $T$ mutation types for each of the $K$ signatures

$\boldsymbol{E}_{N,K}$ : Exposure matrix, containing the contribution of each of the $K$ signatures to each of the $N$ samples

$\boldsymbol{W}_{K,O}$ : Coefficients of logistic regression, weighing contribution of each of the $K$ exposures to prediction of each of the $O$ classes

with:

$N$ : number of samples (and mutational profiles)

$T$ : number of mutation types

$K$ : number of signatures

$O$ : number of output classes, corresponding to DNA repair pathway deficiencies and control

#### A.2. Loss function and update rules for optimization by gradient descent

The loss function $\mathcal{L}_{tot}$ optimized by S-NMF combines a reconstruction loss $\mathcal{L}_r$ with a classification loss $\mathcal{L}_c$ weighed by a factor $\lambda_c$ as follows.

$$\mathcal{L}_{tot} = \mathcal{L}_r + \lambda_c \mathcal{L}_c$$

The reconstruction loss $\mathcal{L}_r$ is defined as the Frobenius reconstruction error of the decomposition of matrix $\boldsymbol{X}$ into matrices $\boldsymbol{E}$ and $\boldsymbol{S}$.

$$\mathcal{L}_r = ||\boldsymbol{X} - \boldsymbol{E}\boldsymbol{S}||_F^2$$

The classification loss $\mathcal{L}_c$ is the categorical cross-entropy loss:

$$\mathcal{L}_c = -\sum_{n=1}^{N}\sum_{o=1}^{O} y_{n,o} \log\left(\hat{y}_{n,o}\right),$$

where $n$ and $o$ are sample and output class indices, respectively, $y_{n,o}$ is the true class label (one-hot encoded), and $\hat{y}_{n,o}$ is the predicted soft class label calculated as follows.

$$\hat{y}_{n,o} = softmax\left(\boldsymbol{E_{n,*}W_{*,o}}\right) = \frac{e^{\boldsymbol{E}_{n,*}\boldsymbol{W}_{*,o}}}{\sum_{p=1}^{P} e^{\boldsymbol{E}_{n,*}\boldsymbol{W}_{*,p}}}$$

The loss function $\mathcal{L}_{tot}$ is optimized using gradient descent, following the iterative updates below.

$$\boldsymbol{S} \leftarrow \boldsymbol{S} - \eta_S \cdot \nabla_{\boldsymbol{S}}\mathcal{L}_{tot}$$
$$\boldsymbol{E} \leftarrow \boldsymbol{E} - \eta_E \cdot \nabla_{\boldsymbol{E}}\mathcal{L}_{tot}$$
$$\boldsymbol{W} \leftarrow \boldsymbol{W} - \eta_W \cdot \nabla_{\boldsymbol{W}}\mathcal{L}_{tot}$$

Symbols $\eta$ denote the respective learning rates (see A.7), and $\nabla$ the (partial-) derivatives of the total loss. In the next sections, we work out the derivatives for the update rules with respect to the reconstruction and the classification losses.

#### A.3. Derivatives of the reconstruction loss:

Firstly, we calculate the derivative with respect to the reconstruction loss. This is identical to what existing methods that applied gradient descent to optimize NMF Lee and Seung (2000). For completeness, we will show the calculation.

Firstly, the frobenius reconstruction loss can be rewritten into four trace terms.

$$\begin{aligned}
||\boldsymbol{X} - \boldsymbol{ES}||_F^2 &= Tr((\boldsymbol{X}^T - \boldsymbol{S}^T\boldsymbol{E}^T)(\boldsymbol{X} - \boldsymbol{ES})) \\
&= Tr(\boldsymbol{X}^T\boldsymbol{X}) - Tr(\boldsymbol{X}^T\boldsymbol{ES}) - Tr(\boldsymbol{S}^T\boldsymbol{E}^T\boldsymbol{X}) + Tr(\boldsymbol{S}^T\boldsymbol{E}^T\boldsymbol{ES}) \\
since, & \\
||\boldsymbol{X}||_F^2 &= Tr(\boldsymbol{X}^T\boldsymbol{X}) \\
Tr(A + B) &= Tr(A) + Tr(B)
\end{aligned}$$

We take the derivative of each of the four trace terms in the loss function. Shown are the derivatives with respect to the exposure matrix $\boldsymbol{E}$, a similar procedure is followed to obtain derivatives with respect to $\boldsymbol{S}$ (applied mathematical rules between parentheses):

$$\begin{aligned}
\nabla_S Tr(\boldsymbol{X}^T\boldsymbol{X}) &= 0 && - \\
\nabla_S Tr(\boldsymbol{X}^T\boldsymbol{ES}) &= \boldsymbol{E}^T\boldsymbol{X} && (\nabla_x Tr(\boldsymbol{AX}) = \boldsymbol{A}^T) \\
\nabla_S Tr(\boldsymbol{E}^T\boldsymbol{S}^T\boldsymbol{X}) &= \boldsymbol{E}^T\boldsymbol{X} && (\nabla_x Tr(\boldsymbol{X}^T\boldsymbol{A}) = \boldsymbol{A}) \\
\nabla_S Tr(\boldsymbol{S}^T\boldsymbol{E}^T\boldsymbol{ES}) &= ((\boldsymbol{E}^T\boldsymbol{E}) + (\boldsymbol{E}^T\boldsymbol{E})^T)\boldsymbol{S} = 2\boldsymbol{E}^T\boldsymbol{ES} && (\nabla_x Tr(\boldsymbol{X}^T\boldsymbol{AX}) = (\boldsymbol{A} + \boldsymbol{A}^T)\boldsymbol{X})
\end{aligned}$$

The final derivatives $\nabla_{\boldsymbol{E}}\mathcal{L}_r$ and $\nabla_{\boldsymbol{S}}\mathcal{L}_r$ of the reconstruction loss $\mathcal{L}_r$ with respect to $\boldsymbol{E}$ and $\boldsymbol{S}$ are as follows.

$$\nabla_{\boldsymbol{S}}\mathcal{L}_r = -2\boldsymbol{E}^T\boldsymbol{X} + 2\boldsymbol{E}^T\boldsymbol{ES} \tag{1}$$

$$\nabla_{\boldsymbol{E}}\mathcal{L}_r = -2\boldsymbol{XS}^T + 2\boldsymbol{ESS}^T \tag{2}$$

#### A.4. Derivatives of the classification loss:

Next, we calculate the gradients of classification component of the total loss. The main difference with a regular logistic regression is that in our case we not only apply gradient descent on the classification weights $W$ but also the exposures $E$ which could considered the input data in a regular logistic regression.

Here we focus on the categorical cross-entropy term and in section A.5 the gradient of the L2-regularization term will be calculated.

$$\mathcal{L}_c = -\sum_{n=1}^{N}\sum_{o=1}^{O} y_{n,o}\log(\hat{y}_{n,o}) + \lambda_{L2}\sum_{w\in\boldsymbol{W}} w^2 \tag{3}$$

For a more convenient calculation of the derivatives, we define $\boldsymbol{Z}$ as the product of the exposures and weights (eq. 5). $\boldsymbol{Z}$ can then be used as input to the softmax.

$$\text{with,} \qquad \hat{y}_{n,o} = softmax(z_{n,o}) = \frac{e^{z_{n,o}}}{\sum_{p=1}^{P} e^{z_{n,p}}} \tag{4}$$

$$\text{with,} \qquad \boldsymbol{Z} = \boldsymbol{EW} \tag{5}$$

Having defined $\boldsymbol{Z}$, we can define the derivative of the cross-entropy loss $d\mathcal{L}_c$ (eq. 6). Taking the derivative using matrices is more complex compared to scalars since the order of the matrices is relevant. Taking this into account, the following rule can be applied:

$$d\mathcal{L}_c = \frac{\partial\mathcal{L}_c}{\partial\boldsymbol{Z}} : dZ \tag{6}$$

$$\text{with,} \qquad \boldsymbol{A} : \boldsymbol{B} = \langle\boldsymbol{A},\boldsymbol{B}\rangle_F$$

where : indicates the Frobenius inner product.

Next, both terms in the derivative of the cross entropy loss $d\mathcal{L}_c$ (eq. 6) need to be calculated.

Firstly, the partial derivative of the cross-entropy loss w.r.t. $\boldsymbol{Z}$ ($\frac{\partial\mathcal{L}_c}{\partial\boldsymbol{Z}}$, eq. 7). This partial derivative is similar to regular logistic regressions, and is as calculated as follow:

$$\frac{\partial\mathcal{L}_c}{\partial\boldsymbol{Z}} = (\hat{\boldsymbol{Y}} - \boldsymbol{Y}) \tag{7}$$

The second term in eq. 6, the full derivative of $\boldsymbol{Z}$ ($dZ$), can be further defined in terms of the exposures and weights:

$$\boldsymbol{Z} = \boldsymbol{EW} \tag{8}$$

$$dZ = dE\boldsymbol{W} + \boldsymbol{E}dW \tag{9}$$

Taken together, $\frac{\partial\mathcal{L}_c}{\partial\boldsymbol{Z}}$ (eq. 7) and $dZ$ (eq. 9) can be substituted in the derivative of the cross entropy loss (eq. 6).

$$\begin{aligned}
d\mathcal{L}_c &= \frac{\partial\mathcal{L}_c}{\partial\boldsymbol{Z}} : dZ \\
&= (\hat{\boldsymbol{Y}} - \boldsymbol{Y}) : (dE\boldsymbol{W} + \boldsymbol{E}dW) \\
&= (\hat{\boldsymbol{Y}} - \boldsymbol{Y}) : dE\boldsymbol{W} + (\hat{\boldsymbol{Y}} - \boldsymbol{Y}) : \boldsymbol{E}dW \\
&= (\hat{\boldsymbol{Y}} - \boldsymbol{Y})\boldsymbol{W}^T : dE + \boldsymbol{E}^T(\hat{\boldsymbol{Y}} - \boldsymbol{Y}) : dW
\end{aligned}$$

Finally, for the gradient w.r.t $\boldsymbol{E}$, $\boldsymbol{W}$ is constant (i.e. $dW = 0$) and for the w.r.t. $\boldsymbol{W}$, $\boldsymbol{E}$ is constant (i.e. $dE = 0$). This result in the final derivatives $\nabla_{\boldsymbol{E}}\mathcal{L}_c$ and $\nabla_{\boldsymbol{W}}\mathcal{L}_c$ of the cross-entropy loss w.r.t. $\boldsymbol{E}$ and $\boldsymbol{W}$:

$$\nabla_{\boldsymbol{W}}\mathcal{L}_c = \frac{\partial\mathcal{L}_c}{\partial\boldsymbol{W}} = \boldsymbol{E}^T(\hat{\boldsymbol{Y}} - \boldsymbol{Y}) \tag{10}$$

$$\nabla_{\boldsymbol{E}}\mathcal{L}_c = \frac{\partial\mathcal{L}_c}{\partial\boldsymbol{E}} = (\hat{\boldsymbol{Y}} - \boldsymbol{Y})\boldsymbol{W}^T \tag{11}$$

#### A.5. Derivative of L2-regularization term

Finally, the derivative of the L2 regularization term w.r.t. $\boldsymbol{W}$. Which is the same as standard derivations.

$$\frac{\partial(\lambda_{L2}\sum_{w\in\boldsymbol{W}} w^2)}{\partial\boldsymbol{W}} = 2\lambda_{L2}W \tag{12}$$

#### A.6. Final gradients

To conclude we combine the partial derivatives of individual terms in the loss function. With regard to the signatures, the derivative only contains a term from of the reconstruction loss (eq. 1). For the exposure, we have a term from both the reconstruction loss (eq. 2) and the cross-entropy loss (eq. 11). For the classifier weight, we have a term from the cross-entropy loss (eq. 10) and from the L2-regularization (eq. 12) of the individual terms in the loss function to get the final gradients of the total loss $\mathcal{L}_{tot}$.

$$\nabla_{\boldsymbol{S}}\mathcal{L}_{tot} = -2\boldsymbol{E}^T\boldsymbol{X} + 2\boldsymbol{E}^T\boldsymbol{ES} \tag{13}$$

$$\nabla_{\boldsymbol{E}}\mathcal{L}_{tot} = -2\boldsymbol{XS}^T + 2\boldsymbol{ESS}^T + \lambda_c(\hat{\boldsymbol{Y}} - \boldsymbol{Y})\boldsymbol{W}^T \tag{14}$$

$$\nabla_{\boldsymbol{W}}\mathcal{L}_{tot} = \lambda_c(\boldsymbol{E}^T(\hat{\boldsymbol{Y}} - \boldsymbol{Y}) + 2\lambda_{L2}\boldsymbol{W}) \tag{15}$$

#### A.7. (Adaptive) Learning rates:

We use the same adaptive learning rates $\eta_S$ and $\eta_E$ for optimization of $\boldsymbol{S}$ and $\boldsymbol{E}$ as in Lee and Seung (2000).

We do not want the integration strength ($\lambda_c$) to effect the optimization of the regression weights $\boldsymbol{W}$, to prevent having a derivative of zero when setting $\lambda_c = 0$. Therefor, we divide our wanted constant learning rate $\mu_W$ by $\lambda_c$. This cancels out the integration strength term ($\lambda_c$) in the derivative $\nabla_{\boldsymbol{W}} \mathcal{L}_{tot}$ (eq.15) and leads to a constant learning rate $\mu_W$ in the final update formula.

$$\eta_S = \frac{\boldsymbol{S}}{2\boldsymbol{E}^T\boldsymbol{E}\boldsymbol{S}} \tag{16}$$

$$\eta_E = \frac{\boldsymbol{E}}{2\boldsymbol{E}\boldsymbol{S}\boldsymbol{S}^T} \tag{17}$$

$$\eta_W = \frac{\mu_W}{\lambda_c} \tag{18}$$

#### A.8. Multiplicative update formulas.

The final multiplicative update formulas are obtained by substituting the derivatives of the loss and the learning rates in the gradient descent update formulas. ($\odot$ and division are element-wise).

$$\boldsymbol{S} \leftarrow \boldsymbol{S} \odot \frac{\boldsymbol{E}^T\boldsymbol{X}}{\boldsymbol{E}^T\boldsymbol{E}\boldsymbol{S}} \tag{19}$$

$$\boldsymbol{E} \leftarrow \boldsymbol{E} \odot \frac{\boldsymbol{X}\boldsymbol{S}^T - \frac{\lambda_c}{2}(\hat{\boldsymbol{Y}} - \boldsymbol{Y})\boldsymbol{W}^T}{\boldsymbol{E}\boldsymbol{S}\boldsymbol{S}^T} \tag{20}$$

$$\boldsymbol{W} \leftarrow \boldsymbol{W} - \mu_W(\boldsymbol{E}^T(\hat{\boldsymbol{Y}} - \boldsymbol{Y}) + 2\lambda_{L2}\boldsymbol{W}) \tag{21}$$

### B. Extended Results

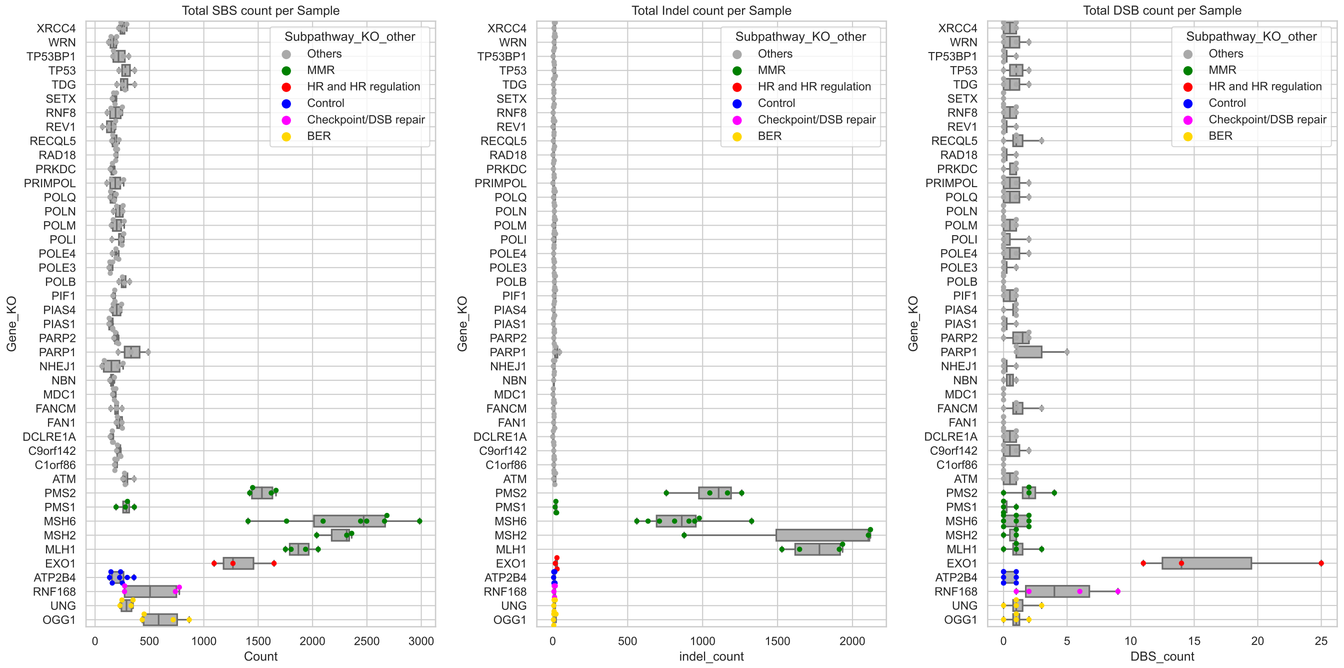


(a) total number of single base substitutions (SBSs) per sample

(b) total number of small insertions and deletions (Indels) per sample

(c) total number of double base substitutions (DBSs) per sample

Fig. S1: **The mutations counts per sample categorized per gene KO**. Grey are the non-distinctive gene KOs. The other samples are the gene KO used for the evaluation of S-NMF colored by the related repair pathway.

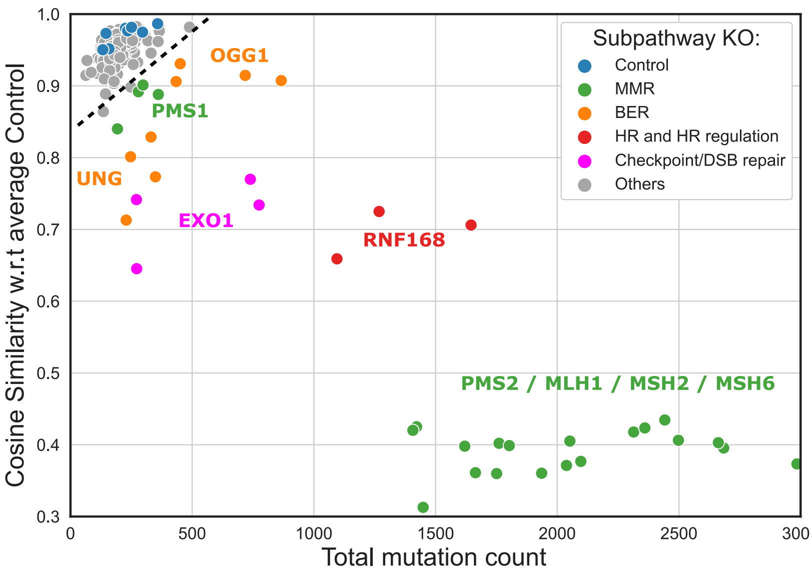


Fig. S2: **Gene knockout and bootstrapped training samples.** Total mutation count and cosine similarity with average profile of control samples, per sample (dot) annotated with gene KO and coloured by repair pathway. The dashed line indicates the threshold used to select mutation profiles: only gene KOs for which all samples yielded higher mutation counts and/or lower similarity to controls than indicated by the dashed line were selected.

| Gene KO | DNA Repair pathway | Total # Samples | # Test | # Train | # Fold 1 | # Fold 2 | # Fold 3 |
|---|---|---|---|---|---|---|---|
| ATP2B4 | Control | 8 | 2 | 6 | 2 | 2 | 2 |
| MSH6 | Mismatch Repair | 8 | 1 | 7 | 3 | 2 | 2 |
| MSH2 | | 3 | 1 | 2 | 0 | 1 | 1 |
| MLH1 | | 4 | 1 | 3 | 1 | 1 | 1 |
| PMS2 | | 4 | 1 | 3 | 1 | 1 | 1 |
| PMS1 | | 4 | 1 | 3 | 1 | 1 | 1 |
| EXO1 | HR | 3 | 1 | 2 | 1 | 0 | 1 |
| RNF168 | | 4 | 1 | 3 | 1 | 1 | 1 |
| OGG1 | Base Exision Repair | 4 | 1 | 3 | 1 | 1 | 1 |
| UNG | | 4 | 1 | 3 | 1 | 1 | 1 |
| | Total: | 46 | 11 | 35 | 12 | 11 | 12 |

Fig. S3: **Table showing the number of samples and how they are subdivided over the test set and training folds.** Shown are the Gene KOs (rows), their annotated DNA repair pathway deficiency and total number of samples/replicates with that gene KO. Next, how the total samples are divided over test and training data, and how the training data is further subdivided into 3 folds for cross-validation.

| **HR** | | **MMR** | **BER** | **NER** | **NHEJ** | **DS** | **DR** | **TLS** | **FA** |
|---|---|---|---|---|---|---|---|---|---|
| BARD1 | BLM | EXO1 | APEX1 | CUL3 | POLL | ATM | ALKBH2 | MAD2L2 | FANCA |
| BRCA1 | BRCA2 | MLH1 | APEX2 | CUL5 | NHEJ1 | ATR | ALKBH3 | POLH | FANCB |
| BRIP1 | EME1 | MLH3 | PARP1 | ERCC1 | XRCC5 | ATRIP | MBD4 | POLK | FANCC |
| GEN1 | MUS81 | MSH2 | POLB | ERCC2 | XRCC6 | CHEK1 | MGMT | POLN | FANCD2 |
| NBN | PALB2 | MSH3 | TDG | ERCC4 | | RNMT | | POLQ | FANCI |
| RAD50 | RAD51 | MSH6 | TDP1 | ERCC6 | | TOPBP1 | | REV1 | FANCL |
| RBBP8 | SHFM1 | PMS1 | UNG | XPA | | TP53 | | REV3L | FANCM |
| TOP3A | TP53BP1 | PMS2 | | XPC | | | | SHPRH | UBE2T |
| XRCC2 | | | | | | | | | |

**Table 2.** List of repair genes included in the analysis and the corresponding pathway annotated. (HR: homologous recombination; MMR: mismatch repair; BER: base excision repair; NER: nucleotide excision repair; NHEJ:non-homologous end joining; DS: damage sensing; DR: direct repair; TLS: translesion synthesis; FA: fanconi anemia).

| **Gene**: | **Recall**: | $Y_{MMRd}$ (avg): | $E_{MMRd}$ (avg): |
|---|---|---|---|
| **MLH1** | 1.0 (30/30) | 1.0 | 1.0 |
| **MLH3** | 1.0 (1/1) | 1.0 | 1.0 |
| **MSH3** | 1.0 (3/3) | 1.0 | 0.63 |
| **MSH2** | 1.0 (2/2) | 1.0 | 0.67 |
| **EXO1** | 1.0 (1/1) | 1.0 | 1.0 |
| **Total** | **1.0** (37/37) | 1.0 | 0.96 |

(a) **HR**

| **Gene**: | **Recall**: | $Y_{HRd}$ (avg): | $E_{HRd}$ (avg): |
|---|---|---|---|
| **BRCA2** | 1.0 (6/6) | 0.95 | 0.75 |
| **BRCA1** | 0.67 (2/3) | 0.49 | 0.51 |
| **RAD50** | 1.0 (1/1) | 1.0 | 0.93 |
| **GEN1** | 1.0 (1/1) | 1.0 | 0.98 |
| **XRCC2** | 1.0 (1/1) | 0.96 | 0.6 |
| **Total** | **0.92** (11/12) | 0.85 | 0.71 |

(b) **MMR**

**Table 3.** Recall of repair pathway deficiency in tumours with homozygous mutated repair gene in the respective pathway.

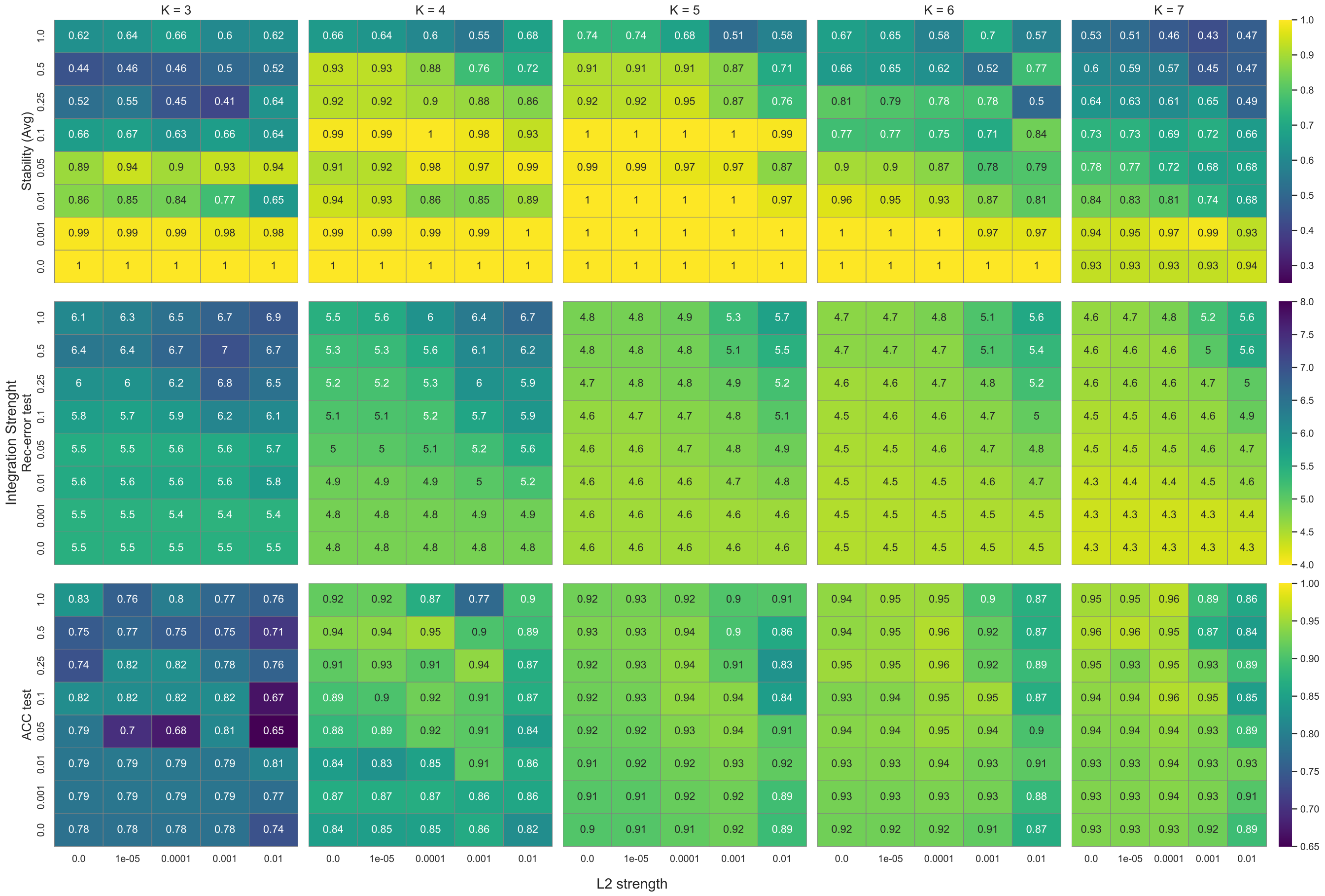


Fig. S4: **Heatmap showing hyperparameter optimization results.** First row) the average stability of the signatures. Second row) reconstruction error. Third row) prediction accuracy. Each cell in the square indicates a the score for that metric of a hyperparameter setting and is annotated and colored according to the the value for the particular metric. The main columns indicated the number of signatures $K$, ranging from 3 to 7. Within each square, the rows indicate the integration strength, and the columns the (L2) regularization strength.

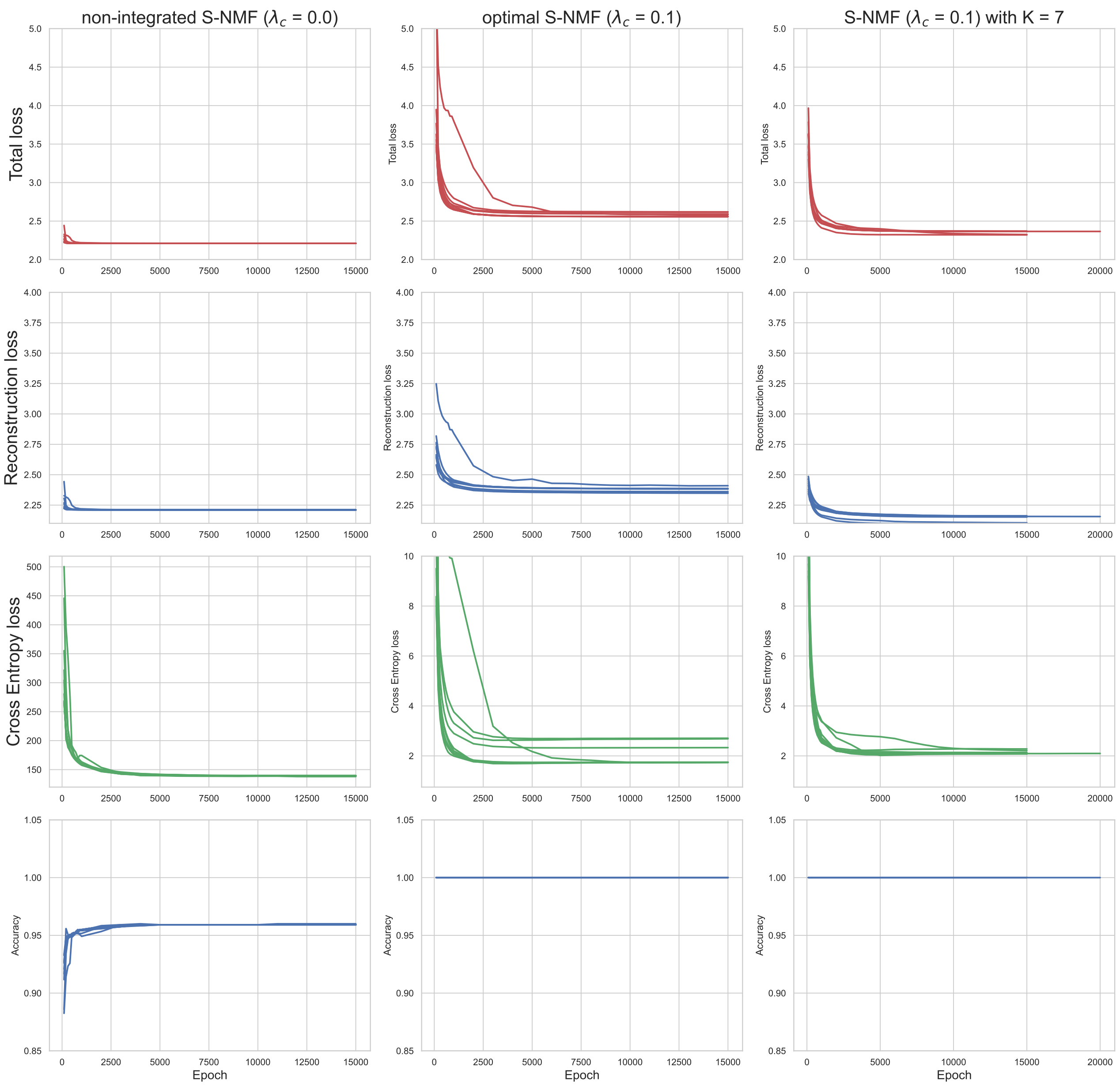


Fig. S5: **Training curves showing development of loss during the training epochs.** Left) non-integrated S-NMF. Middle) optimal S-NMF. Right) S-NMF with K=7. Top row) total loss. 2nd row) Reconstruction loss (frobenius reconstruction error). 3rd row) Cross-entropy loss. Bottom row) Prediction accuracy on training data

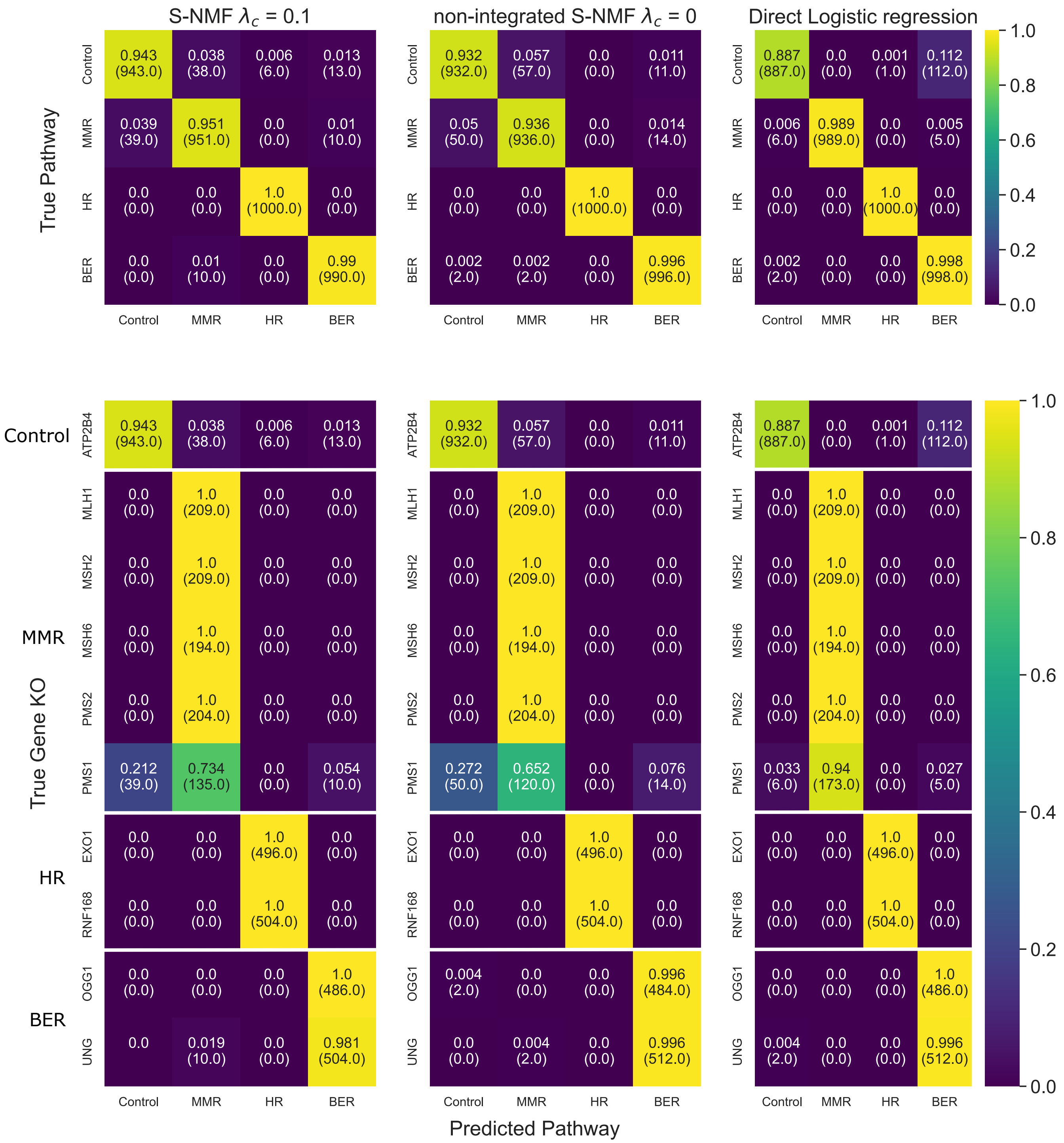


Fig. S6: **Confusion matrices with respect to pathway and gene KO.** Labeled with the percentage of sample that is predicted to the concerned pathway per true pathway/gene KO, the coloring is based on the same percentage. (In parentheses is the total number of sample with the concerned prediction) Overall prediction accuracy for integrated S-NMF is 0.971, non-integrated S-NMF is 0.966, and direct logistic regression 0.968.

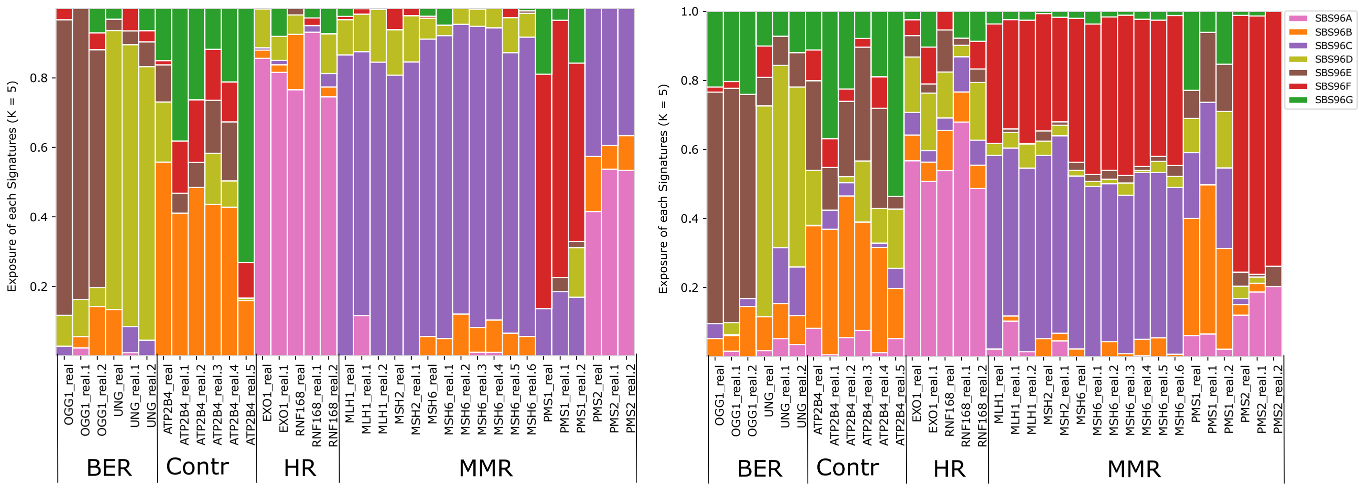


(a) Exposure found by the integrated S-NMF ($\lambda_c = 0.1$) with $K = 7$. (b) Exposure found by the non-integrated S-NMF ($\lambda_c = 0.0$) with $K = 7$.

Fig. S7: **Exposure of the signatures (with $K = 7$) found by S-NMF**. Left) integrated S-NMF. Right) non integrated S-NMF. Only the real samples are shown to allow for practical visualization.

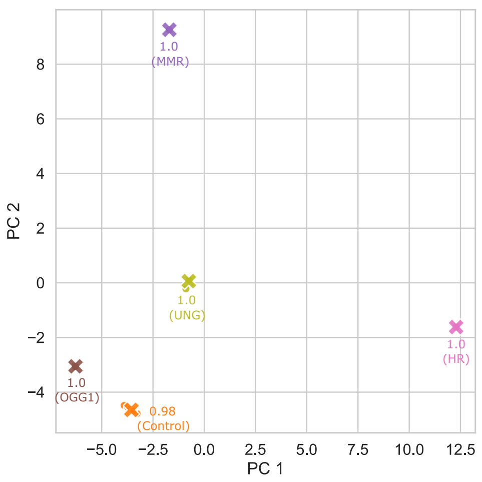


(a) Non-integrated SNMF (average stability = 1.0)

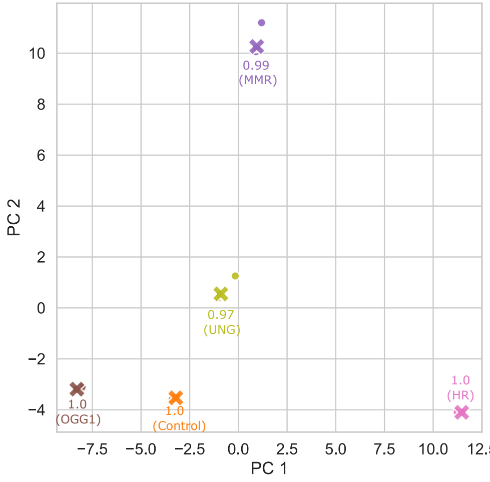


(b) integrated SNMF (average stability = 0.99)

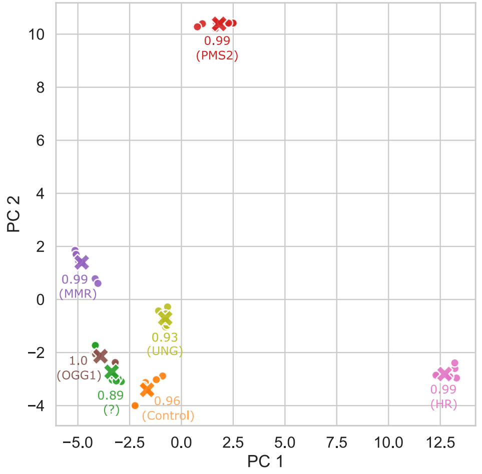


(c) Non-integrated SNMF with K = 7 (average stability = 0.95)

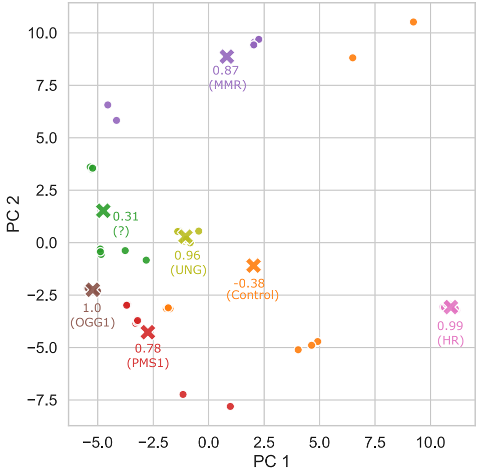


(d) integrated SNMF with K = 7 (average stability = 0.65)

Fig. S8: **PCA plot of signatures of the 10 training runs and the final signatures.** Dots (•) represent the 10 training runs, crosses (×) the centroid of each cluster (i.e. the final signatures). The color indicate the clusters found by partition-clustering (coloring correspond to exposure in Fig. 4). Each centroid is annotated with the stability of the cluster and the repair pathway or gene KO in which the final signature had the highest exposure. Top row) $K = 5$. Bottom row) $K = 7$. Left column) non-integrated S-NF. Right column) integrated SNMF.

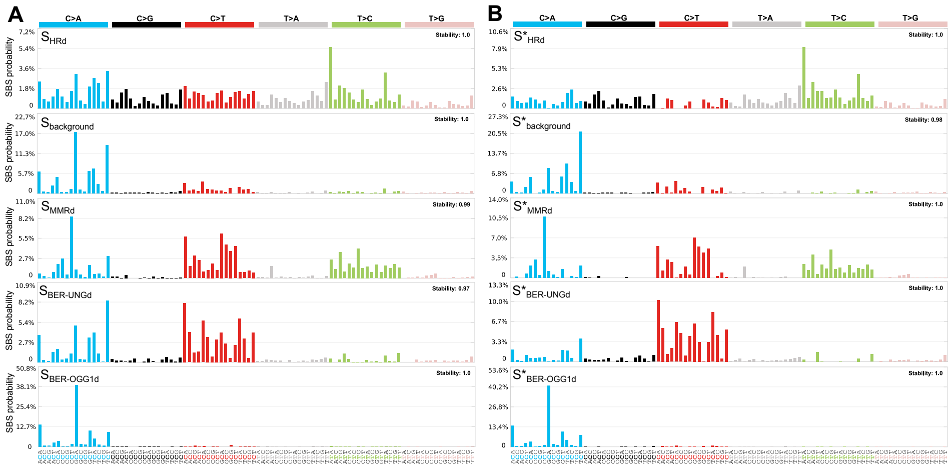


Fig. S9: **Cell line signatures identified by integrated SNMF and NMF (A-B)** Signatures found by (A) integrated SNMF and unsupervised NMF (B) in the cell line samples.

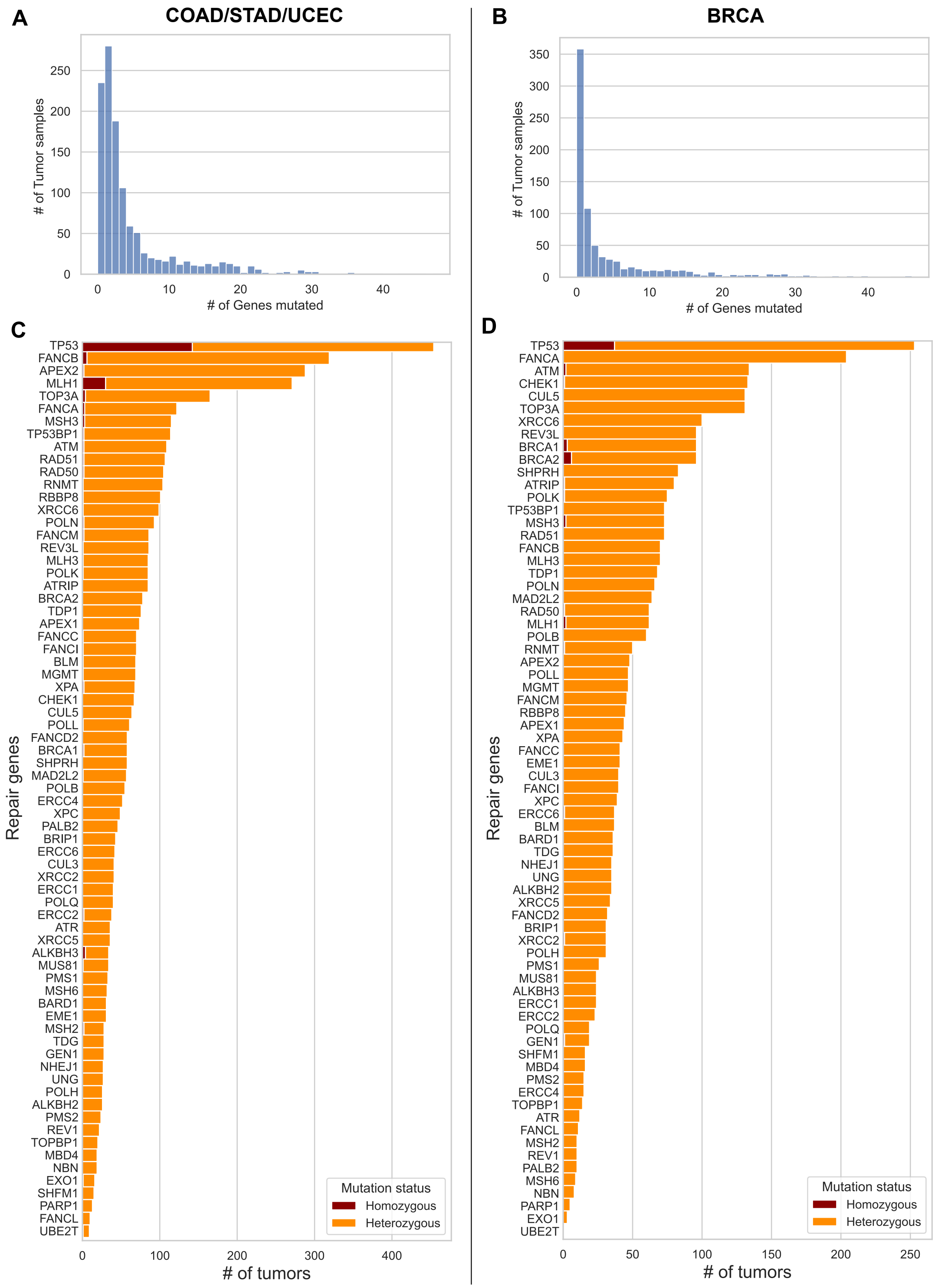


Fig. S10: **Analysis of number of mutated genes. Left)** MMRd related cancer types (in total 1176 tumour samples). **Right)** HRd related cancer types (in total 791 tumour samples). **A and C)** Histogram showing the number of mutated genes. **B and D)** For each gene, the number of tumours that had a (heterozygous or homozygous) loss of function mutation.

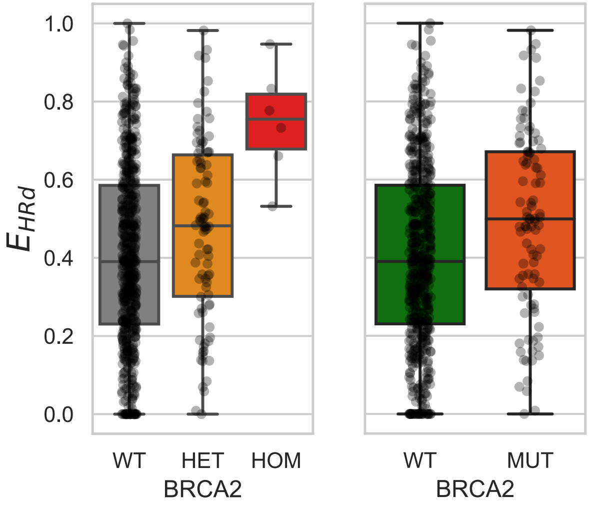


Fig. S11: **Boxplot showing the exposure of the HRd signature for BRCA2 tumours. Left)** All tumours split into three groups based on BRCA2, WT (grey) vs heterozygous (orange) vs homozygous (red) mutated. **Right)** the two groups used in the MW U test where WT has no homozygous mutation in any HR repair gene (green) and combining HET + HOM into single group of mutated tumours (dark orange).

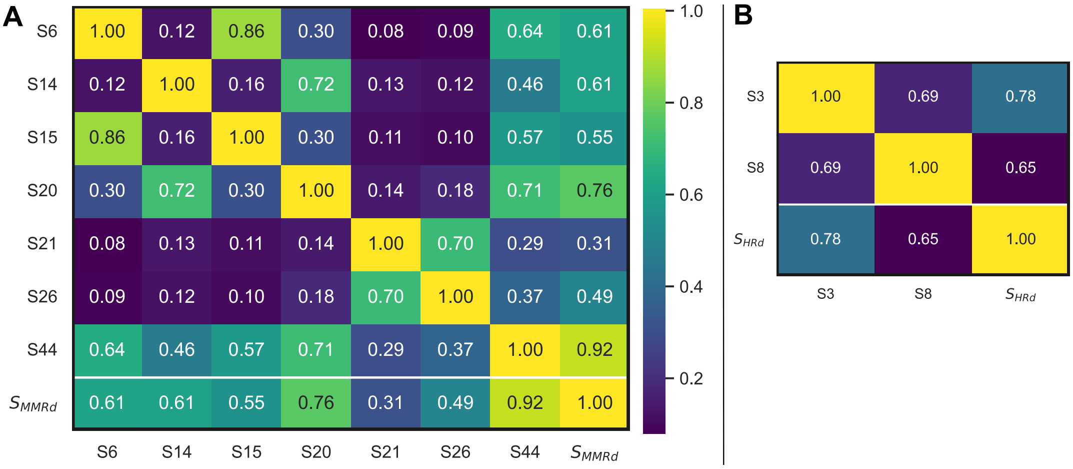


Fig. S12: **Cosine similarity between MMR-related COSMIC signatures ($S$) and the S-NMF signature related to the same repair pathway defi ciency. A)** MMRd related COSMIC signatures and S-NMF MMRd-signature ($S_{MMRd}$). **B)** HRd related COSMIC signatures and S-NMF HRd-signature ($S_{HRd}$).

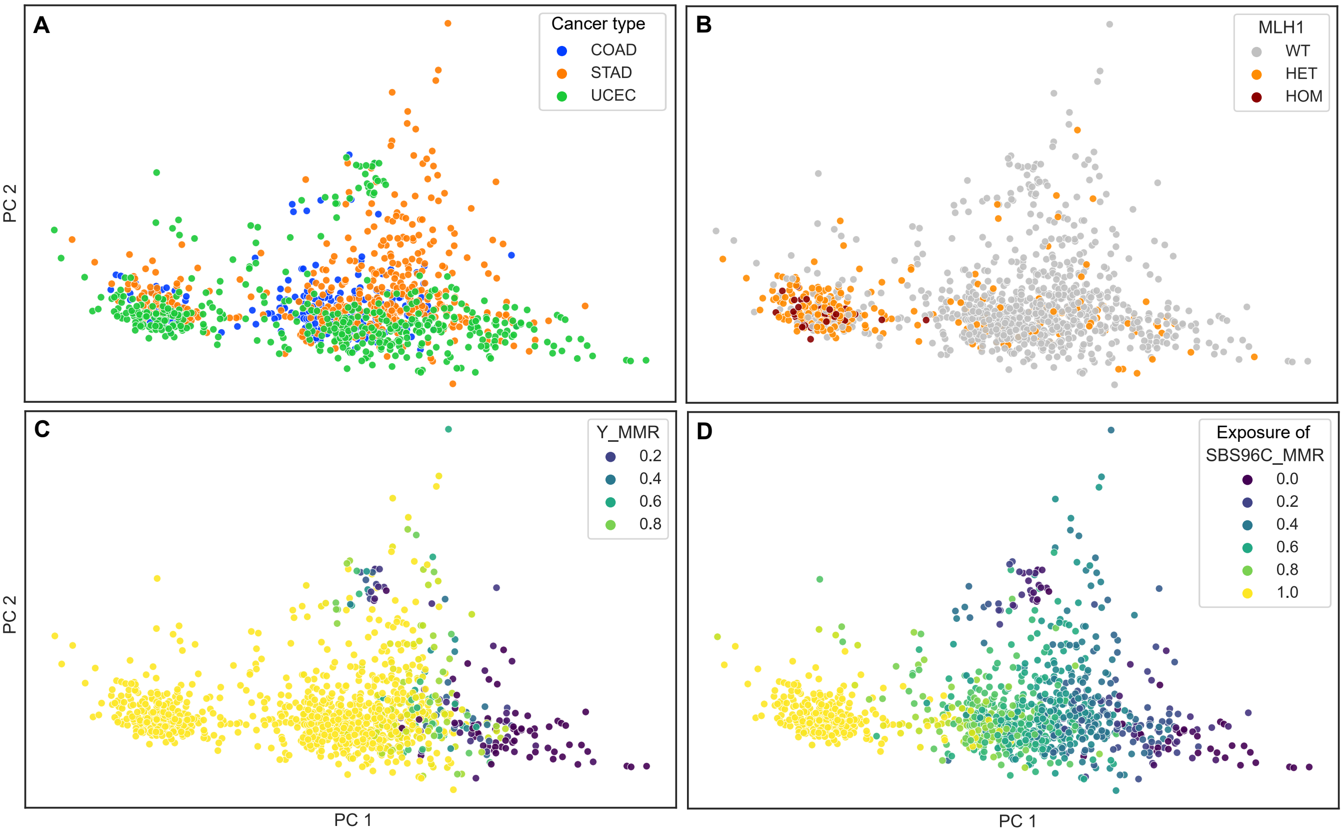


Fig. S13: **PCA of mutational profiles of patients with MMR-related cancer types colon (COAD), stomach (STAD) and uterine (UCEC). A)** the cancer type op each tumour. **B)** the mutation status of MLH1 (WT: wild-type, HET: heterozygous-, HOM: homozygous-mutation). **C)** Prediction probability of MMR-d (Y_MMR). **D)** Exposure of MMR-d related signature (SBS96C_MMR).

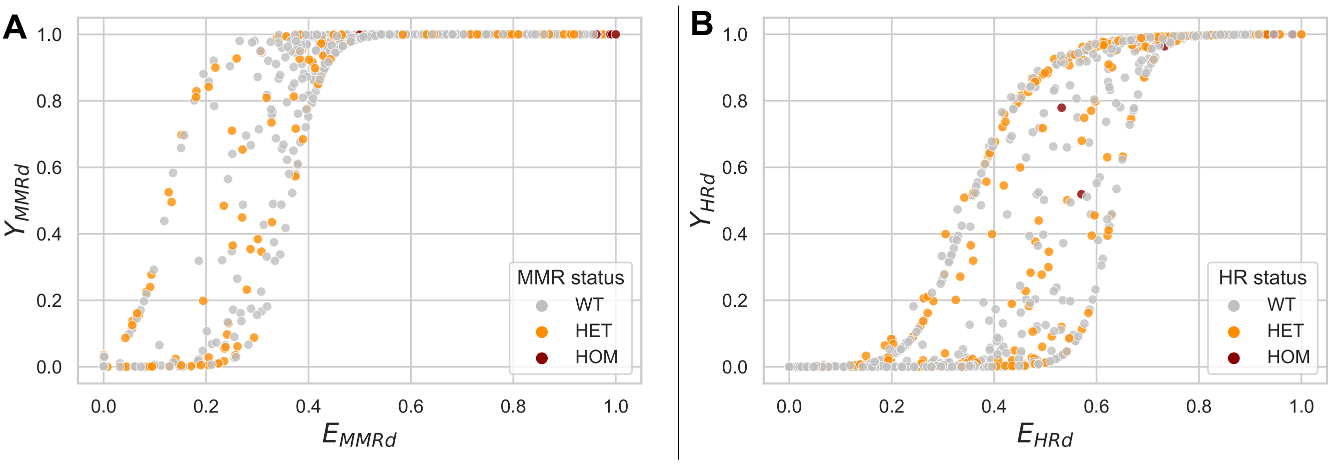


Fig. S14: **Comparing exposure (x-axis) to prediction probability (y-axis) of repair pathway deficiency for all tumours.** Indication of the decision boundary of the SNMF model on the exposure of **A)** MMR-d signature in MMR-related cancer types (COAD/STAD/UCEC) **B)** HR-d signature in HR-related cancer type (BRCA).

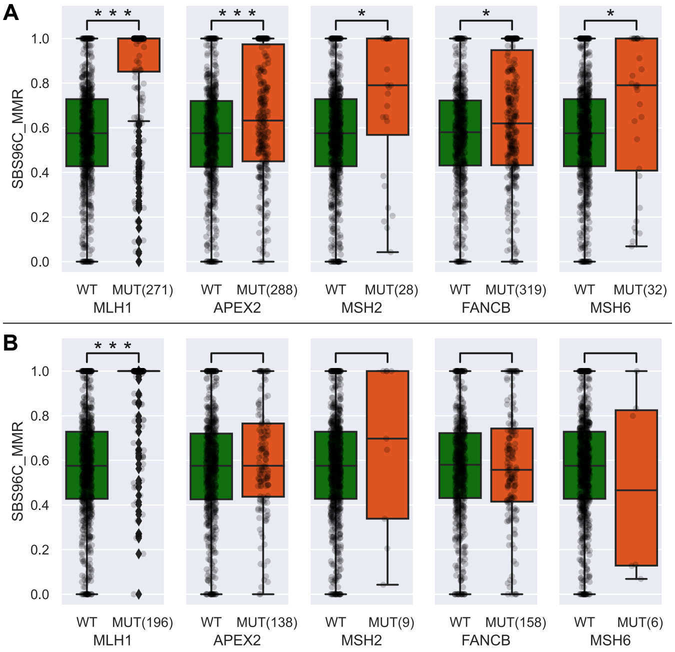


Fig. S15: **Boxplot showing effect of gene on MMRd signature exposure for genes that had a significant association.** Wild-type (WT, green) and mutated samples (MUT, orange, with number of samples): *** $p \leq 0.001$; ** $p \leq 0.01$; * $p \leq 0.05$. **Top)** Original comparison. **Bottom)** Only considering tumours without additional MMR gene mutated.

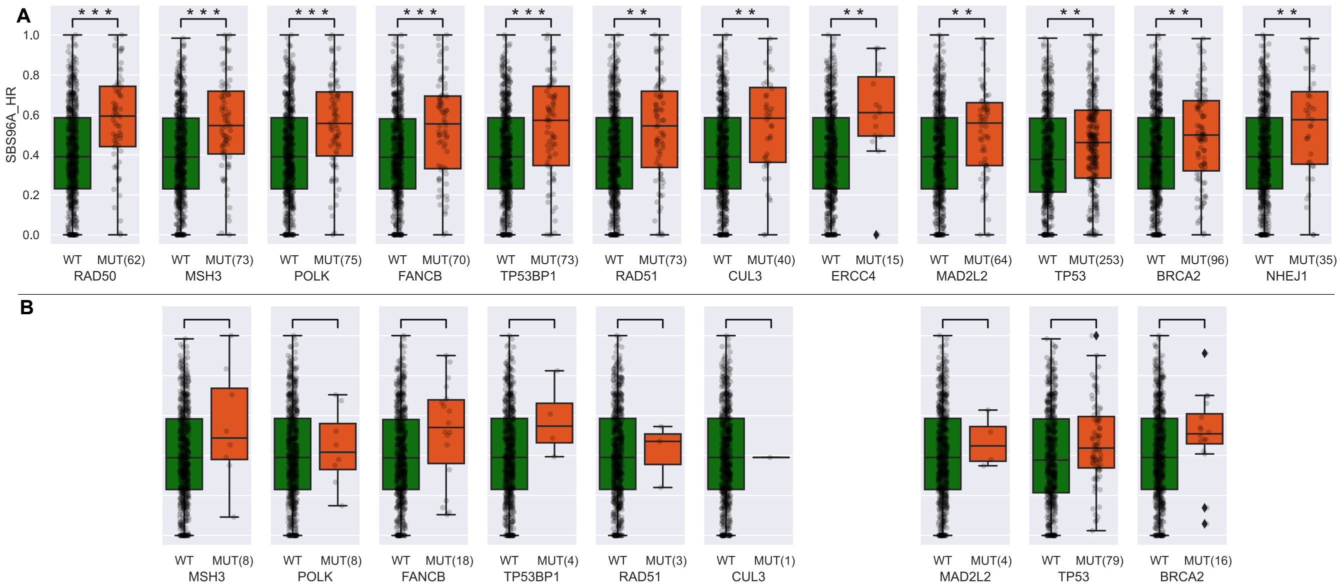


Fig. S16: **Boxplot showing effect of gene on HRd signature exposure for genes that had a significant association.** Wild-type (WT, green) and mutated samples (MUT, orange, with number of samples): *** $p \leq 0.001$; ** $p \leq 0.01$. **Top)** Original comparison. **Bottom)** Only considering tumours without additional HR gene mutated. Empty spots mean there were no tumour samples that did not have an additional mutated HR gene for that gene.

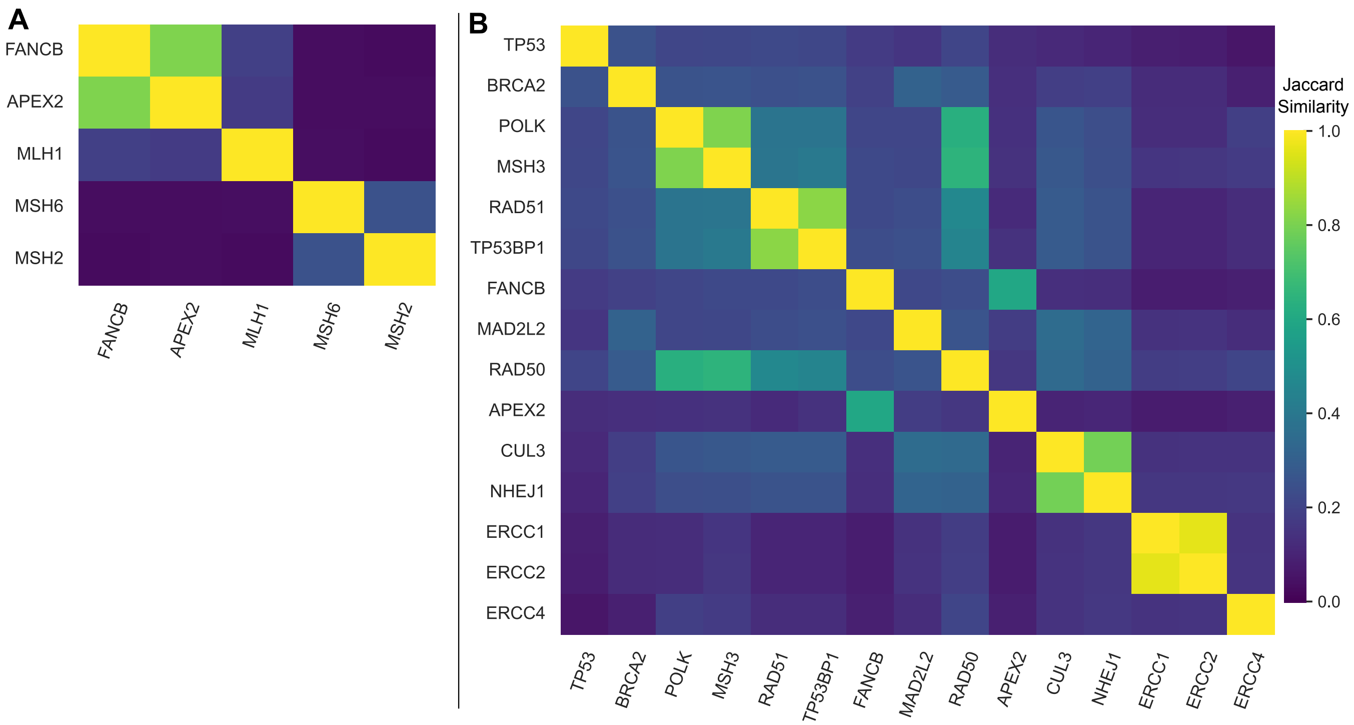


Fig. S17: **Co-occurence (measured by Jaccard Similarity (JS)) between the genes that had a significant effect on the respective pathway deficiency.**. **A)** Co-occurrence (JS) of MMRd associated genes in COAD/STAD/UCEC tumour samples. **B)** Co-occurrence (JS) of HRd associated genes in BRCA tumour samples.
